# *Wolbachia* and host intrinsic reproductive barriers contribute additively to post-mating isolation in spider mites

**DOI:** 10.1101/2020.06.29.178699

**Authors:** Miguel A. Cruz, Sara Magalhães, Élio Sucena, Flore Zélé

**Affiliations:** Centre for Ecology, Evolution and Environmental Changes (cE3c), Faculdade de Ciências da Universidade de Lisboa, Edificio C2, 3º Piso Campo Grande, 1749-016 Lisboa, Portugal; Departamento de Biologia Animal, Faculdade de Ciências da Universidade de Lisboa, Campo Grande, Edifício C2, 1749-016 Lisboa, Portugal; Instituto Gulbenkian de Ciência, Apartado 14, 2781-901 Oeiras, Portugal

**Keywords:** Reproductive manipulation, reproductive isolation, reproductive interference, hybridization, speciation, haplodiploidy

## Abstract

*Wolbachia* are widespread maternally-inherited bacteria suggested to play a role in arthropod host speciation through induction of cytoplasmic incompatibility, but this hypothesis remains controversial. Most studies addressing *Wolbachia*-induced incompatibilities concern closely-related populations, which are intrinsically compatible. Here, we used three populations of two genetically differentiated colour forms of the haplodiploid spider mite *Tetranychus urticae* to dissect the interaction between *Wolbachia*-induced and host-associated incompatibilities, and to assess their relative contribution to post-mating isolation. We found that these two sources of incompatibility act through different mechanisms in an additive fashion. Host-associated incompatibility contributes 1.5 times more than *Wolbachia*-induced incompatibility in reducing hybrid production, the former through an overproduction of haploid sons at the expense of diploid daugters (*ca*. 75% decrease) and the latter by increasing the embryonic mortality of daughters (by *ca*. 49%). Furthermore, regardless of cross direction, we observed near-complete F1 hybrid sterility and complete F2 hybrid breakdown between populations of the two forms, but that *Wolbachia* did not contribute to this outcome. This study identifies the mechanistic independence and additive nature of host-intrinsic and *Wolbachia*-induced sources of isolation. It suggests that *Wolbachia* could drive reproductive isolation in this system, thereby potentially affecting host differentiation and distribution in the field.

## Introduction

In the last decades, it has become increasingly clear that speciation is a continuous process (the “speciation continuum”; Hendry et al. 2000; Powell et al. 2013; Burri et al. 2015; Supple et al. 2015). Ongoing hybridization is taxonomically widespread, and ample variation in the extent and permeability of various reproductive barriers occurs both within and between species (Pinto et al. 1991; Mallet 2008; Hendry et al. 2009; Nosil et al. 2009). Moreover, theoretical studies show that stable partial reproductive isolation can be relatively common (reviewed by Servedio and Hermisson; 2020).

Partial reproductive isolation between lineages (*i*.*e*. differentiated populations or incipient species) can evolve in both sympatry and allopatry due to divergent (including disruptive; Rueffler et al. 2006) sexual and/or ecological selection, and/or as a result of stochastic processes (Schluter 2001, 2009; Turelli et al. 2001; Bolnick and Fitzpatrick 2007; Maan and Seehausen 2011; Nosil 2012). Additionally, in arthropods, partial (or complete) reproductive isolation between and within lineages can result from infection by different cytoplasmically-inherited bacterial reproductive manipulators (Duron et al. 2008; Engelstädter and Hurst 2009), among which *Wolbachia* is the most widespread (Weinert et al. 2015). This endosymbiont can induce various phenotypes of reproductive manipulation in its hosts, including the most common cytoplasmic incompatibility (CI; Werren et al. 2008; Engelstädter and Hurst 2009). CI is a conditional sterility phenotype resulting in increased embryonic mortality of offspring from crosses between infected males and uninfected females (or females harbouring an incompatible strain). Thus, *Wolbachia*-induced CI (wCI) can lead to substantial barriers to gene flow between individuals with different infection status, and could act as an agent of speciation (Laven 1959; Werren 1998; Bordenstein et al. 2001; Telschow et al. 2005; Jaenike et al. 2006). However, whether it plays a significant role in host speciation is still a matter of controversy, mainly because *Wolbachia* can rapidly invade host populations (*i*.*e*. most individuals rapidly become infected, thus immune to CI), and because wCI must produce a sufficient barrier to gene flow to allow nuclear divergence between populations (Hurst and Schilthuizen 1998; Werren 1998; Weeks et al. 2002; Bordenstein 2003). Nevertheless, stable infection polymorphisms are often found in natural populations of many host species (*e*.*g*. Vavre et al. 2002; Keller et al. 2004; Zhang et al. 2013; Hamm et al. 2014; Zélé et al. 2018a). Moreover, whereas speciation solely induced by wCI may require very specific conditions, this *Wolbachia*-induced reproductive manipulation could still play a significant role in host speciation by interacting with other (intrinsic) isolation mechanisms.

The fact that natural populations of many organisms often display variable degrees of reproductive isolation (Scopece et al. 2010; Jennings et al. 2011; Corbett-Detig et al. 2013; Harrison and Larson 2014) offers an excellent opportunity to address the role of wCI in ongoing speciation processes. Still, this has been addressed in a few systems only, and three different, contrasting, scenarios have been described: (1) either no wCI was found in interspecific crosses (Maroja et al. 2008; Gazla and Carracedo 2009; Cooper et al. 2017); (2) *Wolbachia* alone was responsible for post-mating isolation between species through bidirectional wCI (Bordenstein et al. 2001); (3) *Wolbachia* and host genetic factors acted jointly, either in the same direction of crosses (*e*.*g*. a few crosses in Gotoh et al. 2005), or in opposite direction (thereby complementing each other in establishing bidirectional reproductive isolation between species; Shoemaker et al. 1999; Dean and Dobson 2004; see also Gebiola et al. 2016 for CI induced by *Cardinium*). However, when both sources of incompatibility jointly reduce gene flow between genetically differentiated host populations and incipient species, whether they have additive or interacting effects, and precise quantification of their relative contribution to post-mating isolation, has not been addressed. This is at odds with the relevance of such data to better understand the exact contribution of *Wolbachia* to ongoing processes of speciation in arthropods.

*Tetranychus* spider mites constitute an excellent system to address the interplay between symbiont-induced and host intrinsic reproductive incompatibilities. Indeed, they are arrhenotokous haplodiploids (*i*.*e*. males arise from unfertilized eggs and females from fertilized eggs Helle and Bolland 1967), which allows assessing fertilization failure by measuring sex-ratios. Moreover, as many arthropod species, spider mites are often infected with different CI-inducing (or non-inducing) *Wolbachia* strains, whose prevalence greatly varies in natural populations (ranging from 0 to 100%; Gotoh et al. 2003, 2007; Zhang et al. 2016; Zélé et al. 2018a). Due to haplodiploidy (see Breeuwer and Werren 1990; Vavre et al. 2001), wCI can have two different consequences in spider mites, depending on the population tested (*e*.*g*. Gotoh et al. 2003; Perrot-Minnot et al. 2002). In most cases, as in diploid species, eggs from uninfected females fail to hatch when fertilized by sperm from *Wolbachia*-infected males, but wCI affects only the female offspring because males arise from unfertilized eggs (Female mortality - FM-CI type incompatibility; Breeuwer 1997; Vala et al. 2002; Gotoh et al. 2007; Xie et al. 2010; Suh et al. 2015; Bing et al. 2019; Zélé et al. 2020b). In other cases, wCI leads to complete elimination of the paternal set of chromosomes after fertilization of the egg, which successfully develops as a viable haploid male instead of female (Male development - MD-type incompatibility; Vala et al. 2000; Perrot-Minnot et al. 2002; Gotoh et al. 2003). In both cases, the penetrance of wCI (*i*.*e*. the number of embryos affected) greatly varies among populations (from 0 to more than 90% for FM-type and from 0 to 100% for FM-type wCI; Perrot-Minnot et al. 2002; Vala et al. 2002; Gotoh et al. 2007; Xie et al. 2010; Suh et al. 2015; Zélé et al. 2020b), though the origin (*i*.*e. Wolbachia* strain, host genetic background, or both) of such variability in wCI patterns and penetrance is still unknown in spider mites.

Regardless of *Wolbachia* manipulation, variable degrees of reproductive isolation have been found both between and within *Tetranychus* species (*e*.*g*. Keh 1952; Takafuji and Fujimoto 1985; Navajas et al. 2000; Sato et al. 2015; Clemente et al. 2016; Knegt et al. 2017), including between two recently diverged colour forms of the well-studied species *Tetranychus urticae* (Chen et al. 2014; Matsuda et al. 2018). These two closely-related forms have a worldwide distribution (Migeon and Dorkeld 2020), they share the same host plant range (Auger et al. 2013), and they can even be found on the same individual plant (Lu et al. 2017; Zélé et al. 2018a). Therefore, they naturally co-occur and possibly often interact in the field (but see Blanchet et al. 2020). Due to complete reproductive isolation among some populations of the two forms, they were historically described as separate species (*T. urticae* and *T. cinnabarinus*, for the ‘green’ and the ‘red’ form, respectively; Boudreaux 1956; Van de Bund and Helle 1960; Helle and Van de Bund 1962; Smith 1975). Nevertheless, due to morphological and biological synonymy (Auger et al. 2013), and given that many populations of the two forms are not fully reproductively isolated (Murtaugh and Wrensch 1978; Dupont 1979; de Boer 1982b,a; Sugasawa et al. 2002), subsequent studies reclassified them as semi-species (Goka et al. 1996) or members of the same species (Dupont 1979; Fry 1989; Gotoh et al. 1993; Auger et al. 2013). Taken together, these studies thus suggest that speciation is currently ongoing in this species complex, but the role played by wCI in such process is as yet unknown. Indeed, almost all studies addressing reproductive isolation in this system pre-date the identification of *Wolbachia* in spider mites by Breeuwer and Jacobs (1996), and, to our knowledge, only two studies have been conducted since then. One of these showed partial incompatibility (interbreeding was performed for 5 generations) between a *Wolbachia*-uninfected red-form population and a green-form population infected by a non-CI-inducing strain (Sugasawa et al 2002). The other study showed full reproductive isolation between one green-form population and two red-form populations, but *Wolbachia* infection was not assessed (Lu et al 2017).

Here, we assessed the interplay and the relative contribution of wCI and host-associated incompatibilities (HI) on post-mating isolation between three naturally *Wolbachia*-infected populations, two from the red form and one from the green form of *T. urticae*. We performed all possible crosses between *Wolbachia*-infected and *Wolbachia*-free populations in a full-factorial design and measured the embryonic and juvenile mortality of the offspring, as well as the proportion of males and females produced from each cross, over two generations.

## Methods

### Spider mite populations

Three different populations of spider mites, all collected in Portugal and naturally infected with *Wolbachia*, were used in this study. Two populations, ‘Ri1’ and ‘Ri2’, belong to the red form of *T. urticae* and share the same *ITS2* rDNA and *COI* mtDNA sequences. The third population, ‘Gi’, belongs to the green form of *T. urticae* and differs from the former two populations in both *ITS2* rDNA and *COI* mtDNA (*cf*. detailed information in Box S1). The *Wolbachia* strains infecting Ri1 and Ri2 are mutually compatible but induce different levels of cytoplasmic incompatibility despite identical MLST profiles (Zélé et al. 2020b). The *Wolbachia* strain infecting Gi, however, slightly differs from the former two based on MLST and whether it induces CI in this population was heretofore unknown. Since field collection (*cf*. Box S1), these populations were reared in the laboratory under standard conditions (24±2°C, 16/8h L/D) at very high numbers (*ca*. 500-1000 females per population) in insect-proof cages containing bean plants (*Phaseolus vulgaris*, cv. Contender seedlings obtained from Germisem, Oliveira do Hospital, Portugal).

### Antibiotic treatments

After collection, subsets of Gi, Ri1 and Ri2 populations were treated with antibiotics to obtain the corresponding *Wolbachia*-free populations Gu, Ru1 and Ru2. For logistic reasons, the populations Gu and Ru2 used in each of the two experiments reported here were created from two different antibiotic treatments. For Experiment 1, Gu was obtained from a treatment performed in November 2013, and Ru1 and Ru2 from treatments performed in February 2014. Briefly, 100 Gi and 30 Ri1 or Ri2 adult females were installed in petri dishes containing bean leaf fragments, placed on cotton soaked in a tetracycline solution (0.1%, w/v) for three successive generations (Breeuwer 1997; Zélé et al. 2020b). For Experiment 2, Ru1 came from the previous antibiotic treatment but Gu and Ru2 were obtained from new treatments performed in September 2016 and January 2017, respectively. In this case, 300 Gi or Ri2 adult females were installed in petri dishes containing fragments of bean leaves placed on cotton soaked in a rifampicin solution (0.05%, w/v) for one generation (Gotoh et al. 2005; Zélé et al. 2020a). All antibiotic treatments were performed in the same standard conditions as population rearing (24±2°C, 16/8h L/D). After treatment, *Wolbachia*-free populations were maintained without antibiotics in the same mass-rearing conditions as the *Wolbachia*-infected populations for a minimum of three generations to avoid potential side effects of antibiotics (Ballard and Melvin 2007; Zeh et al. 2012; O’Shea and Singh 2015). Subsequently, pools of 100 females from each population were checked by multiplex PCR as described by Zélé et al. (2018b) to confirm their *Wolbachia* infection status before performing the experiments.

### Experiment 1: F1 production and viability

The combined effect of *Wolbachia*- and host-associated incompatibilities (wCI and HI, respectively) on offspring production was investigated by performing all crosses between *Wolbachia*-infected and uninfected individuals from all populations in a full factorial design. These crosses were organized into 5 different categories, each with a different purpose (*cf*. Table 1).

**Table 1.** Description of the five categories of crosses performed in this study.

| Category | Type of crosses | Crosses (♀ x ♂) |
| --- | --- | --- |
| 1 - Controls | intra-population crosses using ♀ and ♂ with the same infection status | Ru1 x Ru1 and Ri1 x Ri1<br>Ru2 x Ru2 and Ri2 x Ri2<br>Gu x Gu and Gi x Gi |
| 2 – Test for wCI only | intra-population crosses using uninfected ♀ and infected ♂ | Ru1 x Ri1<br>Ru2 x Ri2<br>Gu x Gi |
| 3 – Test for HI only (without *Wolbachia*) | inter-population crosses using uninfected ♀ and uninfected ♂ | Ru1 x Ru2 or Gu<br>Ru2 x Ru1 or Gu<br>Gu x Ru1 or Ru2 |
| 4 – Test for wCI-HI interaction | inter-population crosses using (un)infected ♀ and infected ♂ | Ru1 or Ri1 x Ri2 or Gi<br>Ru2 or Ri2 x Ri1 or Gi<br>Gu or Gi x Ri1 or Ri2 |
| 5 – Test for HI only (with *Wolbachia*, to verify that infection itself, in absence of wCI, does not affect HI)* | inter-population crosses using infected ♀ and uninfected ♂ (incl. intra-population controls) | Ri1 x Ru2 or Gu<br>Ri2 x Ru1 or Gu<br>Gi x Ru1 or Ru2<br>(Ri1 x Ru1, Ri2 x Ru2, Gi x Gu) |

\*crosses not performed simultaneously with the others in Experiment 1. The corresponding results were thus analysed separately (cf. Box S2) and are presented in the supplementary materials (Table S1; Figures S1 and S2).

Ten days prior to the onset of the experiment (day −10), age cohorts were created for each infected and uninfected population, by allowing 3*100 mated females (*i*.*e*. ‘female cohorts’) and 4*25 virgin females (*i*.*e*. ‘male cohorts’) to lay eggs during 3 days on detached bean leaves placed on water-soaked cotton. Eight days later (day −2), female nymphs undergoing their last moulting stage (‘quiescent females’ hereafter) were randomly collected from each female cohort and placed separately on bean leaf fragments (*ca*. 9 cm^2^) to obtain virgin adult females with similar age. Virgin males used in the experiment were directly obtained from the male cohorts. On the first day of the experiment (day 0), 1 virgin female and 1 virgin male were installed together on 2.5 cm^2^ bean leaf discs for 3 days before being discarded (day 3). The number of unhatched eggs was counted 5 days later (day 8), and the numbers of dead juveniles, adult males and females were counted 12 days later (day 15).

The experiment was conducted in a growth chamber with standard conditions (24±2°C, 60% RH, 16/8 h L/D). All types of crosses were performed simultaneously, each with 50 independent replicates distributed within two experimental blocks performed one day apart (*i*.*e*. 25 replicates per block). However, given the high number of possible types of crosses (*i*.*e*. 36 combinations) and associated workload, the crosses of category 5 were performed *ca*. 23 months later with minor differences in the methodology (*cf*. details in Box S2). Therefore, data obtained with this latter category were analysed separately and are provided in the supplementary materials (Table S1, Figures S1 and S2).

To calculate the overproduction of F1 males in the brood (MD-type incompatibility; *e*.*g*. Breeuwer and Werren 1990; Navajas et al. 2000; Vala et al. 2000; Vavre et al. 2001) or embryonic mortality of fertilized offspring (*i*.*e*. only females in haplodiploids, hence FM-type incompatibility; Vavre et al. 2000; Vala et al. 2002; Gotoh et al. 2007; Suh et al. 2015; Zélé et al. 2020b), we used indexes adapted from Poinsot et al (1998; see also Cattel et al. 2018; Zélé et al. 2020). MD-type incompatibility was computed as the proportion of sons produced in each cross relative to the control crosses:

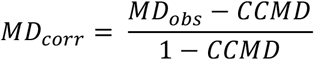

where *MD*_*obs*_ = number of F1 males/total number of eggs, and *CCMD* (calculated as *MD*_*obs*_) is the mean proportion of F1 males observed in control crosses (*i*.*e*. between uninfected individuals of the same maternal population). *MD*_*corr*_ thus takes a value close to 0 when the proportion of males in a given type of cross is similar to that of the controls, but it increases when there is an excess of male production (*i*.*e*. it equals 1 when only sons are produced).

Similarly, FM-type incompatibility was computed as the proportion of embryonic death of daughters produced in each cross relative to the control crosses (hence accounting for variation in background embryonic mortality of both F1 males and females):

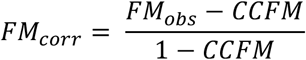

where *FM*_*obs*_ = number of unhatched eggs/[number of unhatched eggs + number of F1 females], and *CCFM* (calculated as *FM*_*obs*_) is the mean embryonic mortality observed in the control crosses. To avoid biases arising from very low numbers of F1 females produced in some inter-population crosses due to MD-type incompatibilities (*cf*. above and results), all females that produced less than two daughters were removed from statistical analyses of *FM*_*corr*_ (*cf*. final sample sizes in Table S1).

Subsequently, to control for potential incompatibilities affecting juvenile viability, we estimated the proportion of dead juveniles in the brood accounting for variation in background juvenile mortality (hence including juvenile mortality of both F1 males and females):

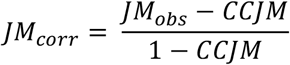

where *JM*_*obs*_ = number of dead juveniles/total number of eggs, and *CCJM* (calculated as *JM*_*obs*_) is the mean juvenile mortality observed in control crosses.

Finally, as both FM- and MD-type incompatibilities affect the proportion of F1 (hybrid) females, their combined effect was determined by assessing the proportion of F1 females resulting from each type of cross:

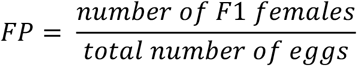

To determine the interplay between FM- and MD-type incompatibilities on hybrid production, we predicted the proportion of F1 females that should be produced in each cross affected by both incompatibilities, assuming that their effects are independent (*H*_0_ hypothesis):

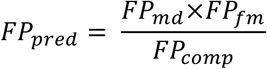

where *FP*_*comp*_ *FP*_*md*_ and *FP*_*fm*_ are, respectively, the mean proportions of F1 females observed in compatible crosses, in crosses affected only by MD-type incompatibility, and in crosses affected only by FM-type incompatibility. Thus, this formula assumes that the decrease in female production due to FM-type incompatibility in crosses already affected by MD-type incompatibility is the same as that the decrease in female production observed between compatible crosses and crosses affected by FM-type incompatibility only (and inversely for MD-type incompatibility). Deviations from this prediction indicate that the two types of incompatibility interfere with each other, that is, they are not independent.

To compare, in each cross affected by both incompatibilities, the observed and predicted proportions of F1 females, we used a Goodness-of-fit Test, with the Pearson goodness-of-fit statistic calculated as follows:

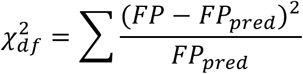

P-values were calculated as the proportion of times the observed proportions of F1 females were equal to or lower than the predicted proportions (Fragata et al. 2014):

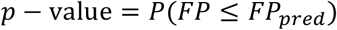

Significant *p*-values thus indicate an interaction between FM- and MD-type incompatibilities, while non-significant *p*-values indicate an independent effect of both types of incompatibility on the proportion of F1 hybrids produced.

### Experiment 2: F1 fertility and F2 viability

To assess the fertility of F1 offspring obtained from inter-population crosses and potential unviability of F2 offspring (*i*.*e*. hybrid breakdown; de Boer 1982b; Sugasawa et al. 2002), all crosses performed in Experiment 1, except those involving Ru2 and Ri2 (because they yielded results similar to Ru1 and Ri1), were repeated in panmixia to obtain large numbers of individuals 13 days prior to the onset of the experiment (day −13). For each cross, 100 virgin females were placed with 100 males (obtained from age cohorts as described for Experiment 1) on an entire bean leaf to produce F1 offspring of the same age. These offspring were used separately to test for F1 female fertility and viability of their offspring (test 1 below) and F1 male fertility and viability of their offspring (test 2 below).

The experiment was conducted in a growth chamber with standard conditions (24±2°C, 16/8 h L/D). In the first test, F1 females from all types of cross were tested simultaneously within four experimental blocks (with a maximum of 25 females per cross tested in each block), while in the second test, uninfected and infected F1 males (*i*.*e*. sons of uninfected or infected females, respectively, independently of the male mated with these females) were tested (and thus analysed) separately. Uninfected F1 males were tested within 3 experimental blocks (with a maximum of 30 males per cross tested in each block); and infected F1 males within 2 experimental blocks (with a maximum of 24 males per cross tested in each block). The number of replicates in each test was limited to the number of F1 offspring that could be obtained from the crosses performed in panmixia (*cf*. final sample sizes in Table S2).

#### Test 1: F1 female fertility and F2 viability

Quiescent F1 females were collected from each cross performed in panmixia and installed on 9 cm^2^ bean leaf fragments 2 days prior to the beginning of experiment (day −2) to emerge as adults while remaining virgin. They were then isolated on 2.5 cm^2^ bean leaf discs on the first experimental day (day 0), and allowed to lay eggs for 4 days, after which they were discarded and the number of eggs laid was counted (day 4). The number of unhatched eggs was counted 5 days later (day 9), and the numbers of dead juveniles and adult males were counted 12 days later (day 16; as mothers were virgin, they could only produce sons).

As F1 female fertility corresponds to their ability to lay a normal number of eggs (Navajas et al. 2000), we estimated both the proportion of ovipositing females and the daily oviposition of these females, taking into account their daily mortality (*i*.*e*. total number of eggs laid by each female/total number of days each female was alive). Hybrid breakdown was assessed as male embryonic and juvenile mortality accounting for variation in background mortality (*i*.*e*. not related to hybrid breakdown). The corresponding *mEM*_*corr*_ and *mJM*_*corr*_ indexes were calculated as follows:

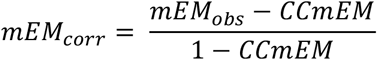

where *mEM*_*obs*_ = number of unhatched eggs/total number of eggs, and *CCmEM* (calculated as *mEM*_*obs*_) is the mean embryonic mortality observed in control crosses (*i*.*e*. category 1);

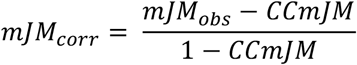

where *mJM*_*obs*_ = number of dead juveniles/total number of eggs, and *CCmJM* (calculated as *mJM*_*obs*_) is the mean juvenile mortality observed in control crosses (*i*.*e*. category 1).

#### Test 2: F1 male fertility and F2 viability

As, in haplodiploids, sons are hemiclones of their mothers, they inherit a single chromosome from each maternal chromosome pair. Thus, in absence of reproductive anomalies they should be fully compatible with females from their maternal population, whereas the expression of an incompatibility may indicate that these males are aneuploid. To test this, adult F1 males were collected from each cross performed in panmixia and placed on 9 cm^2^ bean leaf fragments 2 days prior to the beginning of experiment (day −2) to avoid sperm depletion. On the first experimental day (day 0), each male was installed with one virgin female (obtained from age cohorts created as in Experiment 1) from the same population as its mother on a 2.5 cm^2^ bean leaf disc. Four days were given for the individuals to mate and for the females to lay eggs before both males and females were discarded (day 4). The number of unhatched eggs was counted 5 days later (day 9), and the numbers of dead juveniles, adult males and adult females were counted 12 days later (day 16).

As F1 male fertility corresponds to their ability to sire a normal proportion of offspring (*i*.*e*. F2 females), we estimated both the proportion of males siring daughter(s) and the sex ratio (SR; here calculated as the ratio of females to males because haploid males only sire daughters) in the adult offspring of the females they mated with. Hybrid breakdown was assessed as F2 female embryonic and juvenile mortality accounting for variation in background mortality. As above, *fEM*_*corr*_ and *fJM*_*corr*_ indexes were calculated as:

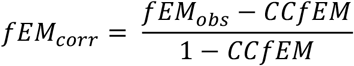

where *fEM*_*obs*_ = number of unhatched eggs/[number of unhatched eggs + number of F2 females] and *CCfEM* (calculated as *fEM*_*obs*_) is the mean embryonic mortality observed in control crosses (*i*.*e*. category 1);

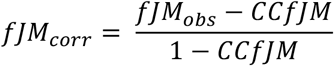

where *fJM*_*obs*_ = number of dead juveniles/[number of dead juveniles + number of F2 females] and *CCfJM* (calculated as *fJM*_*obs*_) is the mean juvenile mortality observed in control crosses (*i*.*e*. category 1).

### Statistical analyses

Analyses were carried out using the R statistical software (v3.6.1). The different statistical models built to analyse the data are described in the Supplementary Materials Table S3. The general procedure to analyse all response variables was as follows: the type of cross was fit as fixed explanatory variable and block was fit as a random explanatory variable. In addition, for the analyses of the proportion of fertile F1 females (*i*.*e*. females that produced at least one egg) and F1 males (*i*.*e*. males that sired at least one daughter), their daily mortality over the 4-day oviposition period was added to the models as it significantly improved their fit. Proportion data were computed as binary response variables (fertile or sterile F1 females and males) or using the function cbind (for female proportion and sex-ratio), except for all corrected variables (*e*.*g*. FM_corr_, MD_corr_, etc.), which are continuous variables bounded between 0 and 1, and for which a “weights” argument was added to the models to account for the number of observations on which they are based. All data were subsequently analysed using generalized linear mixed models with the glmmTMB procedure (glmmTMB package), which allows using a wide range of error distributions that are not implemented in the glmer procedure (Brooks et al. 2017). Proportion data were analysed with a binomial error distribution, or a (zero-inflated) betabinomial error distribution to account for overdispersed errors, and F1 female daily oviposition in experiment 2 was analysed using a log-linked Gaussian error distribution. For all analyses, the significance of the explanatory variable ‘cross’ was established using chi-square tests (Bolker et al. 2009) with the Anova function (car package; Fox and Weisberg 2019). When this explanatory variable was found to be significant, *a posteriori* contrasts were carried out between crosses by aggregating factor levels together and testing the fit of the simplified model using ANOVA (Crawley 2007). Holm-Bonferroni corrections were applied to account for multiple testing (*i*.*e*. classical chi-square Wald test for testing the global hypothesis *H*_*0*_; Holm 1979).

## Results

### F1 offspring production and viability

Reciprocal crosses between naturally *Wolbachia*-infected or *Wolbachia*-free populations of the red (Ri1, Ri2, Ru1 and Ru2) and green (Gi and Gu) form of *T. urticae* allowed testing for the effects of wCI only, HI only, and the combined effect of both sources of incompatibility (*cf. Methods* and Table 1). Overall, we found no significant differences in juvenile mortality among crosses (see Figure 1, Tables S1 and S3), but ample variation in embryonic mortality (*i*.*e*. proportion of unhatched eggs) and/or in male production, both leading to an important decrease in female production (Figures 1 and S1). To identify the sources of such variation (*i*.*e*. wCI and/or HI), we determined which crosses were affected by MD-type incompatibilities (male development; *i*.*e*. overproduction of males resulting from fertilization failure and/or paternal genome elimination) and by FM-type incompatibilities (female embryonic mortality resulting from paternal genome fragmentation). Then, we assessed the consequences of the two types of incompatibility for the resulting proportion of F1 hybrids (only females in haplodiploids).

**Figure 1.**
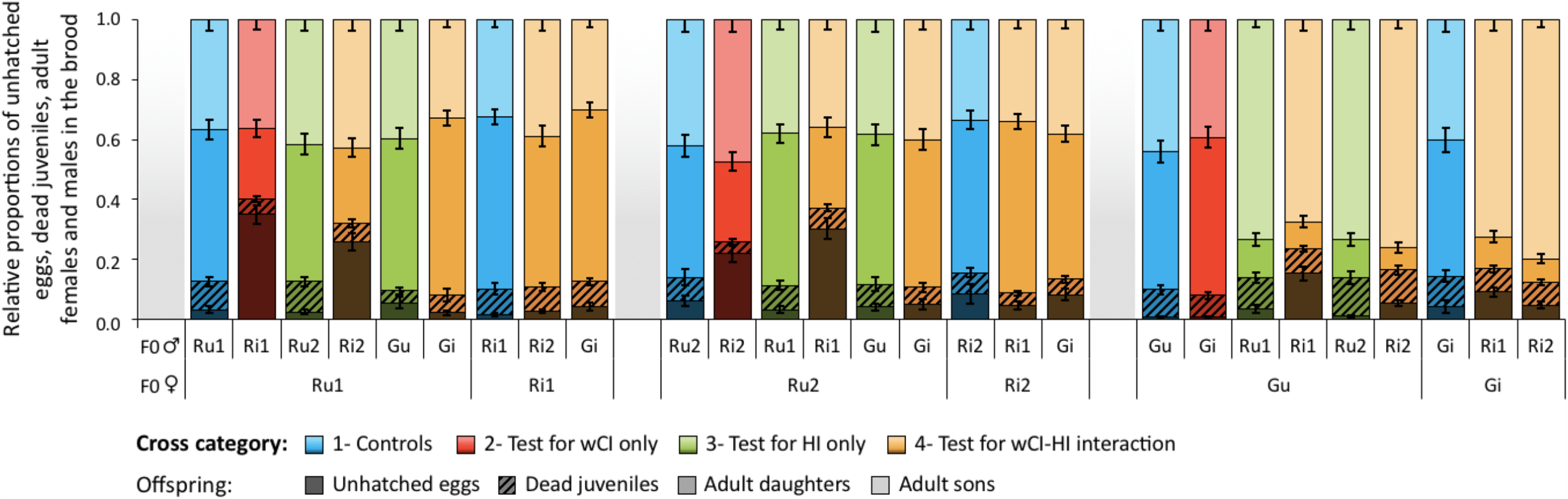
Summary of the development of *T. urticae* eggs resulting from intra- and inter-population crosses between *Wolbachia*-infected and-uninfected mites. Bar plots represent mean ± s.e. relative proportions of unhatched eggs (i.e. embryonic mortality), dead juveniles (i.e. juvenile mortality), adult daughters and sons for each type of cross. Mothers are displayed at the bottom level of the x-axis and fathers on the top level. Note that crosses between infected females and uninfected males (category 5; Figure S1) recapitulate the pattern observed in crosses between uninfected females and uninfected males (categories 1 and 3).

#### Overproduction of males (MD-type incompatibility)

Overall, we found an overproduction of males (*i*.*e*. higher values of the MD_corr_ index; *cf. Methods*) in all inter-population crosses involving females from the green-form population (*ca*. 46.4 to 64.3%) relative to all other crosses (*ca*. 5.6 to 21.5%; *Main cross effect*: χ^2^_26_=460.70, *p*<0.0001; model 1.1, Figure 2a for crosses of categories 1 to 4). Moreover, the level of MD-type incompatibility in these inter-population crosses involving green-form females was not affected by *Wolbachia* infection (*Contrasts among all inter-population crosses using Gu or Gi♀, regardless of* Wolbachia *infection in males*: χ^2^_5_=7.69, *p*=0.17). In contrast, we found no overproduction of males in any of the inter-population crosses involving red-form females (*Contrasts among all crosses with low MD*_*corr*_, *including the controls and regardless of* Wolbachia *infection in both males and females*: χ^2^_20_=26.11, *p*=0.16). Finally, the analysis of crosses involving *Wolbachia*-infected females and uninfected males (*i*.*e*. crosses of category 5; Figure S2a) recapitulated the pattern observed in crosses involving uninfected females and males (*i*.*e*. crosses of categories 1 and 3), further showing that *Wolbachia* infection in females also does not affect MD_corr_. Indeed, as before, higher values of MD_corr_ were found for inter-population crosses involving green-form females (*ca*. 57.9 to 64.5%) as compared to all other crosses (*ca*. 5.9 to 30.3%; *Main cross effect*: χ^2^_8_=174.26, *p*<0.0001; model 1.2; Table S2). Taken together, these results revealed an overproduction of males due to HI between green-form females and red-form males, with *Wolbachia* infection playing no role in this outcome.

**Figure 2.**
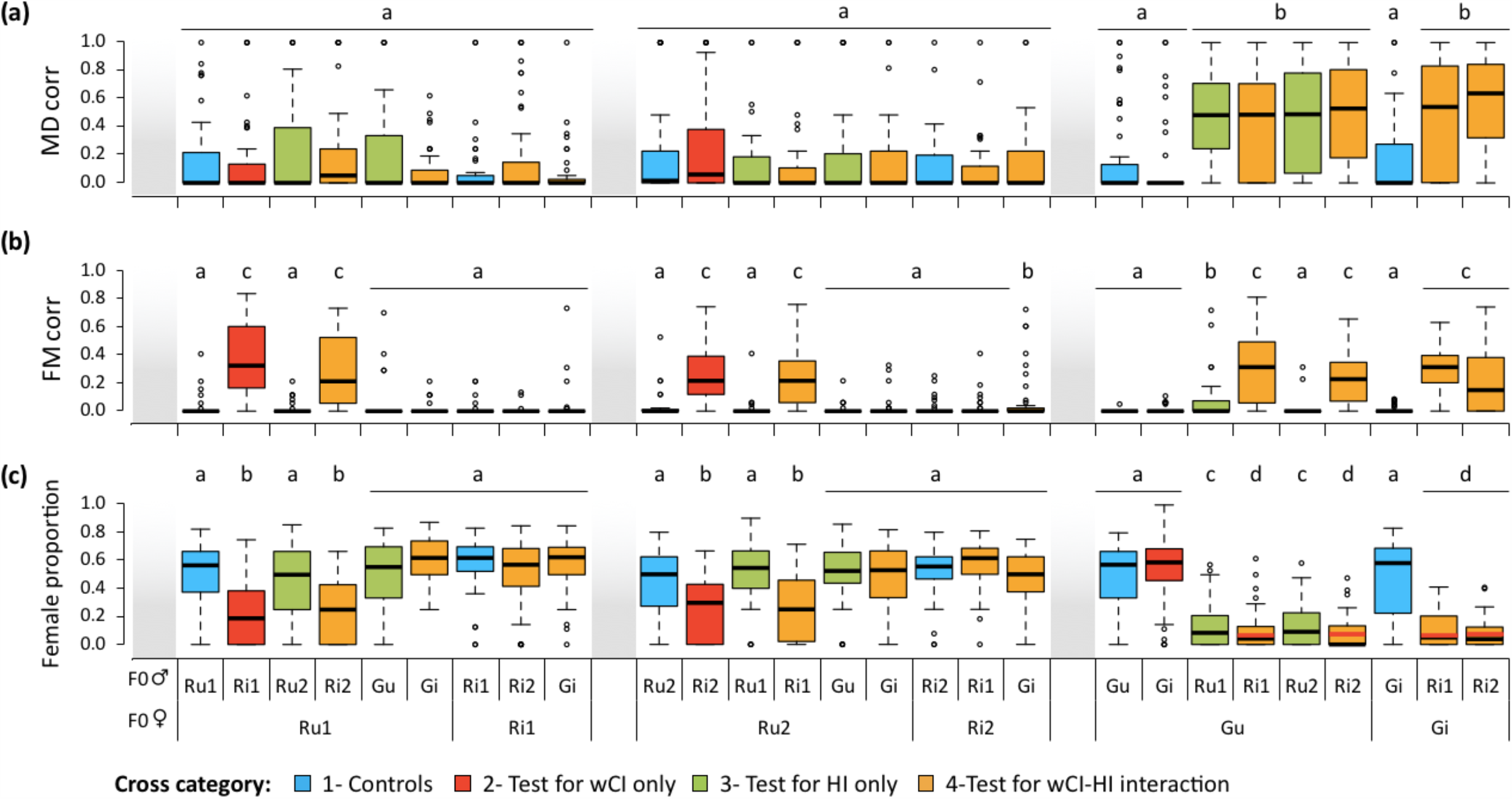
Overproduction of males, female embryonic mortality, and resulting hybrid production in intra- and inter-population crosses using *Wolbachia*-infected and uninfected mites. **(a)** Boxplot of the proportion of males produced in all crosses relative to that in control crosses (MD_corr_). **(b)** Boxplot of the proportion of unhatched eggs relative to females, accounting for the basal level of this proportion observed in control crosses (FM_corr_). **(c)** Proportion of F1 adult females (*i*.*e*. hybrids) in the brood. Horizontal red bars displayed within boxes for crosses affected by both MD- and FM-type incompatibilities indicate predicted values of female proportion (FP_pred_) under the assumption that the two types of incompatibilities have an independent effect on hybrid production. Mothers are displayed on the bottom level of the x-axis and fathers on the top level. Identical or absent superscripts indicate nonsignificant differences at the 5% level among crosses. Note that crosses between infected females and uninfected males (category 5; Figure S2) recapitulate the pattern observed in crosses between uninfected females and uninfected males (categories 1 and 3).

#### Hybrid (female) embryonic mortality (FM-type incompatibility)

Overall, we found higher levels of female embryonic mortality relative to controls (FM_corr_ index; *cf. Methods*) in all crosses using *Wolbachia*-infected red-form males, either crossed with uninfected red-form females (as found by Zélé et al; 2020b), or with green-form females independently of their *Wolbachia* infection status (from 22.2 to 42.7% on average; *Main cross effect*: χ^2^_26_=506.20, *p*<0.0001; model 1.3; Figure 2b). In addition, there were no significant differences among these crosses (χ^2^_7_=8.76, *p*=0.27; despite a tendency for Ri1 males to induce higher levels of FM-type CI than Ri2 males: 35% vs. 29% on average), which shows that the *Wolbachia* strain infecting the green-form population did not rescue (even partially) wCI induced by *Wolbachia* infection in red-form males. All other crosses resulted in no (or low) female embryonic mortality (from 0.2 to 10.5% on average; *Contrasts among all these crosses with low FM*_*corr*_: χ^2^_16_=19.99, *p*=0.22). Thus, these results restrict FM-type incompatibilities between populations to CI induced by *Wolbachia* infection in males from the two red-form populations, with the same penetrance in inter-population and intra-population crosses (hence regardless of HI).

#### Consequences of MD- and FM-type incompatibilities for hybrid (female) production

As a result of the MD- and FM-type incompatibilities described above, we also found ample variation in the proportion of females (FP) produced across crosses (*Main cross effect*: χ^2^_26_=966.45, p<0.0001; model 1.7; Figure 2c). Contrast analyses further revealed four statistically different proportions depending on the type of crosses: (1) *ca*. 51% daughters produced in compatible crosses (*i*.*e*. unaffected by both incompatibilities; *Contrasts among these crosses*: χ^2^_16_=21.22, *p*=0.17); (2) *ca*. 26% daughters produced in crosses affected by FM-type incompatibilities only (*Contrasts among these crosses*: χ^2^_3_=2.98, *p*=0.40; *ca*. 49% decrease compared to compatible crosses: χ^2^_1_=187.67, *p*<0.0001); (3) *ca*. 13% daughters produced in crosses affected by MD-type incompatibilities only (*Contrasts among these crosses*: χ^2^_1_=0.04, *p*=0.84; *ca*. 75% decrease compared to compatible crosses: χ^2^_1_=292.02, *p*<0.0001; and *ca*. 76% decrease when using crosses of category 5: χ^2^_8_=278.23, p<0.0001; model 1.8; Figure S2c); and (4) *ca*. 9% daughters produced in crosses affected by both FM- and MD-type incompatibilities (*Contrasts among these crosses*: χ^2^_3_=3.57, *p*=0.31; *ca*. 82% decrease compared to compatible crosses: χ^2^_1_=606.40, *p*<0.0001).

Both types of incompatibility appeared to have lower consequences on hybrid production when combined than when acting alone. Indeed, we found around 31% decrease in hybrid production due to FM-type incompatibility when comparing groups (3) and (4) (χ^2^_1_=7.49, *p*=0.03) and close to 65% decrease in hybrid production due to MD-type incompatibility when comparing groups (2) and (4) (χ^2^_1_=141.97, *p*<0.0001). However, this was only a consequence of their cumulative effects. Indeed, we found a perfect fit between the observed and the predicted proportions of F1 females for crosses affected by both MD- and FM-type incompatibilities, calculated assuming that both affect hybrid production with the same strength when acting either alone or combined (Figure 2c; Goodness-of-fit test: *Gu♀xRi1♂*: χ^2^_47_=14.30, *p*=0.58; *Gu♀xRi2♂*: χ^2^_47_=8.46, *p*=0.65; *Gi♀xRi1♂*: χ^2^_47_=13.90, *p*=0.56; and *Gi♀xRi2♂*: χ^2^_48_=7.37, *p*=0.59). Thus, these results show that MD- and FM-type incompatibilities, hence HI and wCI (see above), are independent, so that their effects are additive, with the former contributing 1.5 times more in reducing hybrid production than the latter (*ca*. 75% and 49% less hybrids produced, respectively).

### F1 offspring fertility and viability of the F2

To estimate the effects of wCI and HI on the fitness of F1 offspring obtained from all aforementioned crosses (except those involving Ru2 and Ri2 populations, *cf*. Methods), we assessed the fertility of virgin F1 females and of F1 males backcrossed to females from their maternal population, and both embryonic and juvenile mortality of the resulting F2 offspring (*i*.*e*. hybrid breakdown; de Boer 1982b; Sugasawa et al. 2002).

#### Fertility of F1 females and viability of their offspring (Test 1)

The proportion of virgin F1 females that laid at least 1 egg differed significantly depending on the crosses they resulted from (χ^2^_15_=214.26, *p*<0.0001; model 2.1; Figure 3a). While most females resulting from all intra-population crosses oviposited (*ca*. 96% on average; *Contrasts among intra-population crosses*; χ^2^_7_=8.42, *p*=0.30), more than 99% of those resulting from inter-population crosses were unable to lay eggs. Moreover, although 6 hybrid females (over a total of 760), all resulting from crosses between males and females with the same *Wolbachia* infection status (either both infected, or both uninfected), were found to be fertile, they laid very few eggs (average daily oviposition of 0.63 ± 0.15 compared to 6.37 ± 0.09 for females resulting from intra-population crosses; *cf*. Table S3), with no significant difference among inter-population crosses (*Contrasts among inter-population crosses*; χ^2^_7_=8.59, *p*=0.28).

**Figure 3.**
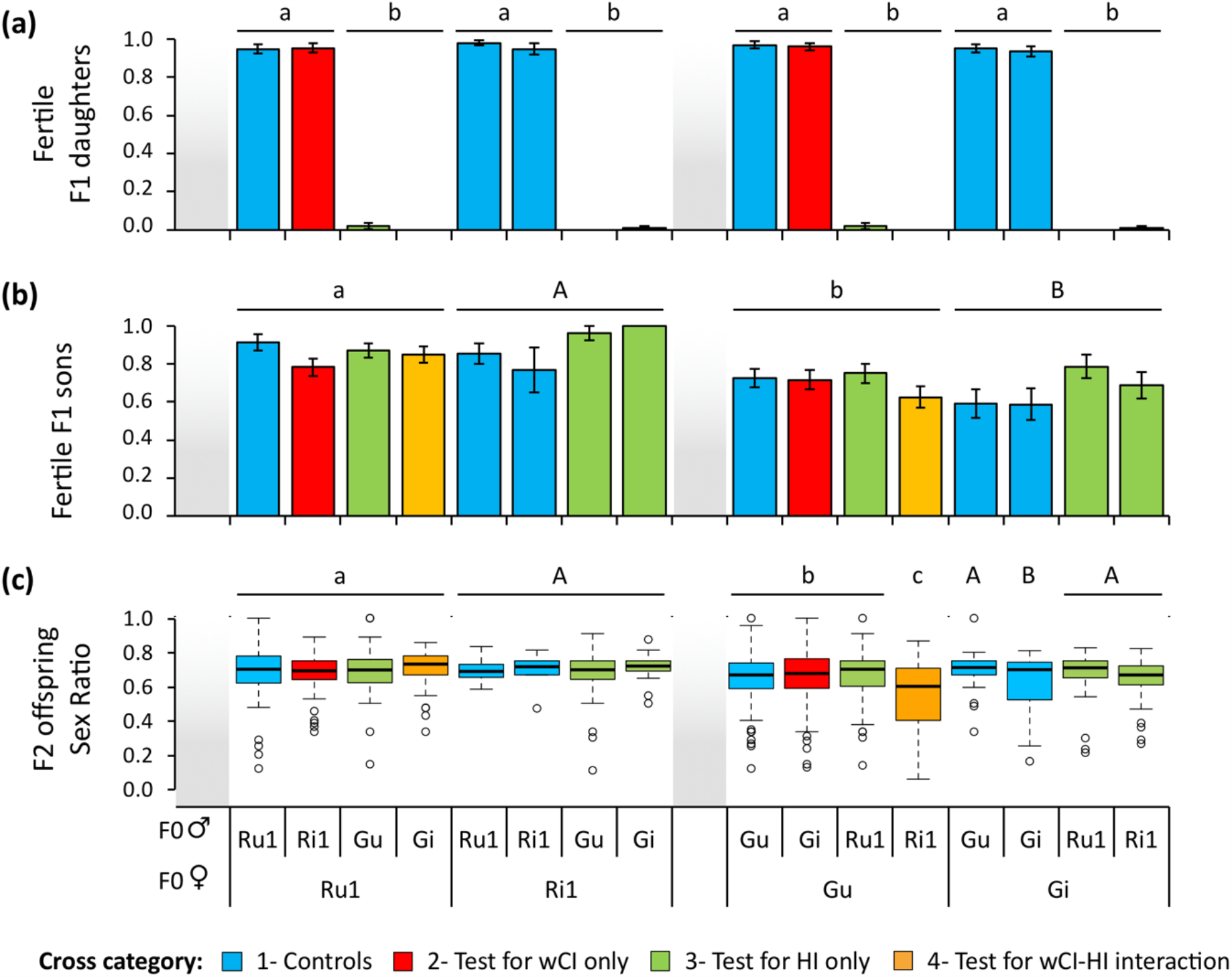
Proportion of fertile F1 female and male offspring resulting from intra- and inter-population crosses using *Wolbachia*-infected and uninfected mites, and sex-ratio of F2 offspring resulting from backcrosses of F1 males. Average proportion (± s.e.) of **(a)** fertile F1 females (*i*.*e*. proportion of females laying at least 1 egg) and **(b)** fertile F1 males (*i*.*e*. proportion of males siring at least 1 daughter when mated with a female from the same population as their mother). **(c)** Boxplot of sex ratio (daughters to sons) of F2 offspring sired by F1 males. Mothers are displayed on the bottom level of the x-axis and fathers on the top level. Identical or absent superscripts indicate nonsignificant differences at the 5% level among crosses.

None of the few eggs laid (15 in total) by the 6 fertile hybrid females hatched (Table S3), which corresponds to full F2 hybrid breakdown. In contrast, embryonic mortality (mEM_corr_) of eggs laid by F1 females resulting from intra-population crosses was very low (*ca*. 5%), with only a very small increased mortality (*ca*. 2%) in the brood of F1 females from the *Wolbachia*-infected green-form population (*Main cross effect:* χ^2^_7_=23.33, *p*=0.001; model 2.3; Figure S3a). Similarly, juvenile mortality (mJM_corr_) in the offspring (*i*.*e*. all F2 males) of virgin F1 females resulting from intra-population crosses was very low (below *ca*. 6%), and varied slightly depending only on their maternal origin (*Main cross effect:* χ^2^_7_=18.57, *p*=0.01; model 2.4; Figure S3b). Indeed, the offspring of infected green F1 females had higher juvenile mortality than the offspring of infected red-form females (independently of their grandfather; *Contrasts between Gi and Ri1 females*: χ^2^_1_=12.53, *p*=0.002), and the offspring of all uninfected F1 females displayed an intermediate mortality (*Contrasts between Gu-Ru1 and Gi females:* χ^2^_1_=4.28, *p*=0.17; *Contrasts between Gu-Ru1 and Ri1 females:* χ^2^_1_=4.49, *p*=0.17). These results thus show that, due to very high hybrid sterility (99% non-ovipositing females) and complete hybrid breakdown, the red- and green-form populations studied here are, in fact, fully post-zygotically isolated (*i*.*e*. no gene flow).

#### Fertility of F1 males and viability of their offspring (Test 2)

The proportion of F1 males siring at least one daughter (when backcrossed with a female from their maternal population) differed significantly depending on the crosses they resulted from (χ^2^_7_=25.58, *p*<0.001; model 2.5.1, and χ^2^_7_=15.23, *p*=0.03; model 2.5.2, for uninfected and infected males, respectively). However, this difference was mainly attributable to the maternal populations of these males and/or to the population of the females they mated with (*i*.*e*. as both are the same, it is not possible to disentangle their effects). Indeed, F1 males mated with (and sons of) green-form females were less fertile than those mated with (and sons of) red-form females (*ca*. 17.39% and 25.97%, for uninfected and infected males, respectively; *cf*. Figure 3b).

The maternal population of fertile F1 males also affected the proportion of daughters they sired, but only when they were uninfected by *Wolbachia* (χ^2^_7_=42.10, *p*<0.0001; model 2.8.1; Figure 3c). In this case, we found that F1 males mated with (and sons of) red-form females sired on average more offspring (*ca*. 69%) than F1 males mated with (and sons of) green-form females (*ca*. 55% for those mated with infected red-form males; χ^2^_1_=32.13, *p*<0.0001; and *ca*. 65% for those mated with all other males; χ^2^_1_=8.96, *p*=0.008). We also found some differences in the sex-ratio of offspring resulting from crosses using infected F1 males (χ^2^_7_=15.19, *p*=0.03; model 2.8.2), but this effect was only due to a higher variance (but not median) in the Gi♀xGi♂ control cross, and no difference was found among all other crosses (χ^2^_6_=9.93, *p*=0.13; Figure 3c).

Finally, neither embryonic mortality (fEM_corr_) nor juvenile mortality (fJM_corr_) varied among the offspring of F1 infected males (*Main cross effect on fEM*_*corr*_: χ^2^_7_=5.58, *p*=0.59; model 2.6.2, and *on fJM*_*corr*_: χ^2^_7_=11.68, *p*=0.11; model 2.7.2). Although both varied among the offspring of F1 uninfected males (*Main cross effect on fEM*_*corr*_: χ^2^_7_=26.31, *p*<0.001; model 2.6.1, *and on fJM*_*corr*_: χ^2^_7_=22.64, *p*=0.002; model 2.7.1), this variation is not attribuable to wCI or HI. Indeed, a higher embryonic mortality (*ca*. 7% on average) was found only in the offspring of uninfected F1 males mated with (and sons of) green-form females compared to those mated with (and sons of) red-form females (*ca*. 4% in average). In line with this, we found that juvenile mortality varied depending on both the maternal and the paternal populations of F1 uninfected males (or the females they mated with), but regardless of incompatibility (due to wCI and/or HI) between the parental populations (see Figure S4b).

Overall, these results indicate that F1 males resulting from all types of incompatible crosses do not suffer a reduction in fertility. This suggests that they are true hemiclones of their mother, thereby escaping both sources of incompatibility (wCI and HI).

## Discussion

Using three populations of the two genetically differentiated colour forms of *T. urticae*, each naturally infected or cured from *Wolbachia*, we assessed the relative contribution of *Wolbachia*-induced CI (wCI) and of host-associated incompatibilities (HI) to post-mating isolation. Our results revealed that both sources of incompatibility jointly reduced the production of F1 hybrid females in the same direction, albeit through different and independent mechanisms, and with HI contributing *ca*. 1.5 times more than wCI to this reduction (*ca*. 75% and 49% less F1 hybrids produced, respectively). Additionally, we found a *Wolbachia*-independent near-complete F1 hybrid female sterility and full F2 hybrid breakdown in all reciprocal crosses between the green- and the red-form populations.

### Expression of host-associated incompatibilities

Crosses performed among uninfected host populations in absence of *Wolbachia* infection confirmed that the two populations belonging to the red form of *T. urticae* were mutually compatible (Zélé et al. 2020b), but were fully isolated from the green-form population. We found three different types of post-mating reproductive barriers between these populations: (1) a sharp and unidirectional (between females from the green-form propulation and males from the red-form populations but not in reciprocal crosses) reduction in F1 hybrid (female) production, concurrent with an increased production of F1 males (*i*.*e*. MD-type incompatibility); (2) near-complete F1 hybrid sterility (> 99%) in all reciprocal crosses between the red and the green-form population; and (3) full F2 hybrid breakdown, as none of the few eggs produced by F1 hybrid females hatched.

MD-type incompatibilities, which result in an excess of F1 males at the expense of daughters, have already been reported between populations from the two colour forms of *T. urticae* (Murtaugh and Wrensch 1978; Sugasawa et al. 2002; Lu et al. 2017), as well as among populations of the same colour form (Navajas et al. 2000; Perrot-Minnot et al. 2004). In haplodiploids, this type of incompatibility can result from either fertilization failure or haploidization of fertilized F1 eggs. Partial haploidization of fertilized eggs is unlikely here, as males surviving such defect would be aneuploid, and thus, should produce fewer daughters, an outcome we did not find when testing F1 males. Moreover, there is yet no evidence that aneuploid embryos can actually be viable in spider mites. Conversely, complete haploidization of fertilized eggs is a plausible explanation, as *Wolbachia* can cause MD-type incompatibility in *T. urticae* (Vala et al. 2000; Perrot-Minnot et al. 2002; Gotoh et al. 2003), and this outcome was shown to result from paternal genome elimination following fertilization in haplodiploids (*i*.*e*. it restores haploidy and thus leads to the production of viable males; Breeuwer and Werren 1990; Tram et al. 2006). However, in spider mites, males are naturally produced from unfertilized eggs (*i*.*e*. arrhenotoky; Helle and Bolland 1967) and not from the elimination of the paternal genome in fertilized eggs (*i*.*e*. pseudo-arrhenotoky; Nelson-Rees et al. 1980; Sabelis and Nagelkerke 1988). Therefore, fertilization failure resulting from a defect at any of the successive stages of the reproductive process in the female reproductive tract is another possible explanation for this type of incompatibility between populations (*e*.*g*. reduction in sperm transfer/storage, sperm ejection/dumping, reduced sperm activation or attraction to the egg, and sperm-egg incompatibility; Zeh and Zeh 1997; see also Takafuji and Fujimoto 1985; Perrot-Minnot et al. 2004). Moreover, although the results presented here do show that premating isolation between the two forms is incomplete (*i*.*e*. no hybrids would be produced in absence of mating), we cannot exclude the possibility that fewer copulations have occurred in these crosses. Only direct observations of copulations, of the fertilization process, and of early embryogenesis of the offspring in these crosses would allow testing these hypotheses.

Irrespective of the underlying mechanisms, we found both asymmetric (MD-type) and symmetric (F1 hybrid sterility and F2 hybrid breakdown) patterns of reproductive incompatibilities between spider mite populations of the two forms. In general, asymmetric incompatibilities have been tied to “Bateson-Dobzhansky-Muller incompatibilities” (BDMIs – negative epistatic interactions between alleles from independently evolving lineages) between autosomal loci and uniparentally inherited factors (*e*.*g*. maternal transcripts; Sawamura 1996; Turelli and Orr 2000; or cytoplasmic elements such as mitochondrial genes; Burton and Barreto 2012). In contrast, symmetrical patterns of incompatibilities are generally associated to BDMIs between nuclear genes inherited from both parents (Turelli and Moyle 2007). This suggests that MD-type incompatibilities are caused by cytonuclear interactions, whereas hybrid sterility and hybrid breakdown are mainly due to incompatibilities between nuclear genes. This is in line with some evidence from previous work using spider mites, albeit several of these studies also highlight a role for cytonuclear interactions in hybrid sterility and hybrid breakdown (Overmeer and van Zon 1976; de Boer 1982b; Fry 1989; Sugasawa et al. 2002; Perrot-Minnot et al. 2004).

### Expression of *Wolbachia*-induced CI within and among populations

Crosses between *Wolbachia*-infected males and uninfected females within and among populations showed that the *Wolbachia* strains infecting the two red-form populations induce imperfect FM-type incompatibility (*ca*. 22 to 43% female embryonic mortality) and are mutually compatible (as found by Zélé et al; 2020b). Here, we further showed that wCI has the same penetrance within and among host populations, including the population from the green form. Conversely, the strain infecting the green-form population did not induce CI within or between populations, neither of the FM-type nor of the MD-type. Moreover, in contrast to some other non CI-inducing *Wolbachia* strains in *T. urticae* (Vala et al. 2002), this strain did not rescue the CI induced by the strain infecting the red-form populations. Taken together, these results show a unidirectional pattern of wCI, which reduces hybrid production between the *Wolbachia*-infected red-form populations and the green-form population, regardless of *Wolbachia* infection in the latter. Finally, we found no evidence for hybrid breakdown (*i*.*e*. increased mortality of F2 offspring produced by F1 females escaping wCI) induced by any of the *Wolbachia* strains, suggesting that such effect is not a common feature in spider mites, or that it is restricted to strains inducing MD-type incompatibilities (Vala et al. 2000).

### The combined effects of incompatibility types for hybrid production and gene flow

In some systems, wCI may play a greater role than HI in reducing gene flow between hosts. For instance, complete post-mating isolation due to bidirectional wCI has been found in interspecific crosses between the mosquitoes *Aedes polyniensis* and *Ae. riversi* (Dean and Dobson 2004), and between the parasitoid wasps *Nasonia giraulti* and *N. vitripennis* (Breeuwer and Werren 1990, 1995), while only partial isolation was found in interspecific crosses upon *Wolbachia* removal (asymmetrical hybrid production and F2 hybrid breakdown, respectively). In other systems, however, CI induced by symbionts and host intrinsic factors can complement each other when acting in opposite directions, as found between *Encarsia gennaroi* and *Cardinium*-infected *E. suzannae* (Gebiola et al. 2016), or can act synergistically to reduce gene flow in the same direction. This was found between some populations of the spider mite *Panonychus mori*, where wCI mainly results in haploidization of fertilized eggs and can increase existing MD-type incompatibilities between populations (Gotoh et al. 2005). However, the relative contribution of wCI and HI to post-mating isolation was not quantified in such cases, let alone whether they have additive or interacting effects.

In our system, we found that HI and wCI act jointly to prevent the production of F1 hybrid offspring in crosses between green-form females and red-form males. Moreover, we showed that they act independently and additively, with HI contributing *c*.*a*. 1.5 times more than wCI to the reduction in F1 hybrid production. However, because all F1 hybrids were either sterile or produced unviable eggs, *Wolbachia* did not affect gene flow between the red- and green-form populations studied here. Nonetheless, these results suggest that wCI may have an important role in restricting gene flow between populations of *T. urticae* that are only partially isolated, which is a common phenomenon in *T. urticae* (*e*.*g*. Dupont 1979; Fry 1989; Navajas et al. 2000; Sugasawa et al. 2002; Perrot-Minnot et al. 2004). In particular, the effects of wCI may be considerable when MD-type incompatibilities between hosts are weaker (Murtaugh and Wrensch 1978; Navajas et al. 2000; Sugasawa et al. 2002), and/or when the two types of incompatibilities act in opposite directions (*e*.*g*. as found in *Cardinium* infected *Encarsia* parasitoid wasps; Gebiola et al. 2016). Therefore, more studies using population pairs with variable degrees of post-mating isolation, and assessing pre-mating isolation, are needed to better understand the extent to which *Wolbachia* can hamper gene flow between natural populations of spider mites, and determine its exact role in the speciation processes currently ongoing in this system.

### Ecological implications of host-associated and *Wolbachia*-induced incompatibilities

Our results show strong reproductive interference (see Gröning and Hochkirch 2008; Burdfield-Steel and Shuker 2011) between the populations from the two forms of *T. urticae* used in our study, which may potentially impact their dynamics by favouring the green-form population. Indeed, green-form females mated with red-form males produce less (sterile) hybrid daughters but more (fertile) sons than red-form females mated with green-form males, and this overproduction of sons may have important consequences for the persistence these populations. Moreover, despite our finding that F1 green-form males had a slightly lower fitness than F1 red-form males (*i*.*e*. lower fertility and higher embryonic mortality of their daughters), their overproduction should allow green-form females to transmit more genes (thereby mitigating the costs of heterospecific matings; Feldhaar et al. 2008). This should also increase, at the next generation, the probability of conspecific matings (*e*.*g*. as in *Callosobruchus* beetles; Kyogoku and Nishida 2012) for green-form females, and of heterospecific matings for red-form females, which may again favour the green-form population.

*Wolbachia* may also affect the balance of the interactions between these populations, both due to the direct effects of infection on host fitness (*i*.*e. Wolbachia* slightly increases the embryonic and juvenile mortality of F2 sons of green-form, but not red-form, F1 females), but also due to wCI. Indeed, although wCI leads to embryonic mortality of hybrid daughters of green-form females, all these daughters are sterile. Conversely, wCI leads to embryonic mortality of fertile daughters of red-form females, which may further disadvantage red-form females in populations that are polymorphic for *Wolbachia* infection (as often found in spider mites; Breeuwer and Jacobs 1996; Zhang et al. 2013; Zélé et al. 2018a). Note, however, that the effect of wCI between partially isolated populations of the two forms (*e*.*g*. de Boer 1982b; Sugasawa et al. 2002) may lead to completely different scenarios, as it could also affect fertile hybrid daughters produced by females of either form.

Such ecological scenarios are likely to occur in natural populations of *T. urticae*, as incompatible populations (both of the same and of different colour forms) often co-occur in the field (Helle and Pieterse 1965; Lu et al. 2017), and the populations used in this study were collected in the same geographical area (*cf*. Box S1). However, these scenarios will also depend on the strength and the symmetry of pre-mating and post-mating prezygotic reproductive barriers between populations (Sato et al. 2015, 2018; Gebiola et al. 2017; Clemente et al. 2018). Indeed, although one study reported no assortative mating between the colour forms of *T. urticae* (Murtaugh and Wrensch 1978), this may vary between populations, as found between *T. urticae* and *T. evansi* (Sato et al. 2014; Clemente et al. 2016). In line with this, contrasting results were found concerning the effect of *Wolbachia* on spider mite mating behaviour (Vala et al. 2004; Rodrigues et al. 2018). Thus, to understand the implications of reproductive interference in this system, future studies should focus on prezygotic isolation between *T. urticae* populations, infected or not by *Wolbachia*.

## Conclusions

Our results show that host-associated and *Wolbachia*-induced incompatibilities in this system lead to different outcomes and that both contribute to counter hybridization between populations of the two *T. urticae* colour forms. Furthermore, these two types of incompatibility have additive effects in the same direction of crosses, which hints at a possible role of *Wolbachia*-induced incompatibilities in host population divergence and subsequent evolution of intrinsic reproductive barriers (*e*.*g*. as found in the *Nasonia* wasps; Bordenstein et al. 2001). Indeed, although the level of divergence between the populations studied here limits our understanding of the contribution by *Wolbachia* in this system (because they are either not or fully isolated), our results suggest that this reproductive manipulator may have a considerable effect between partially isolated populations and, thus, could play an important role in the processes of speciation currently ongoing in spider mites. Finally, our results raise important questions about the ecological consequences of *Wolbachia*-driven reproductive interference in arthropods, and call for further studies to understand its impact on the dynamics and distribution of natural populations from the same species, but also from closely-related species.

## Supporting information

Description of the datasets

Complete dataset Experiment 1

Complete dataset Experiment 2_Test 1

Complete dataset Experiment 2_Test 2

R script Experiment 1

R script Experiment 2_Test 1

R script Experiment 2_Test 2

## Abbreviations

CI: cytoplasmic incompatibility;
wCI: *Wolbachia*-induced cytoplasmic incompatibility;
HI: Host-associated incompatibility;
EM: Embryonic mortality;
FM: Female mortality;
MD: Male development;
JM: Juvenile mortality;
FP: Female proportion over total number of eggs laid;
SR: Sex ratio (here ratio of females to males in the offspring).

## Data accessibility & Supplementary Materials

Full datasets, R scripts and Supplementary Materials (Boxes, Tables and Figures) are available online: https://doi.org/10.1101/2020.06.29.178699.

## Acknowledgements

We are grateful to Salomé Clémente and Leonor Rodrigues for their help in collecting data for the first experiment, to Inês Santos for the maintenance of the spider mite populations and the plants, and to Inês Fragata and Fabrice Vavre for useful discussions and suggestions. This work was funded by an FCT-ANR project (FCT-ANR//BIA-EVF/0013/2012) to SM and Isabelle Olivieri and by an ERC Consolidator Grant (COMPCON, GA 725419) to SM. MC was funded through an FCT PhD fellowship (SFRH/BD/136454/2018), and FZ through an FCT Post-Doc fellowship (SFRH/BPD/125020/2016). Funding agencies did not participate in the design or analysis of experiments. Version 5 of this preprint has been peer-reviewed and recommended by *Peer Community In Evolutionary Biology* (https://doi.org/10.24072/pci.evolbiol.100116).

## Authors’ contributions

Experimental conception and design of the first experiment: FZ with discussions with SM. Experimental conception and design of the second experiment: MC, FZ, SM, ES; Acquisition of data, statistical analyses, and writing of the first version of the manuscript: MC, FZ. Subsequent versions were written with input from SM and ES. All authors have approved the final version for publication.

