## Supplementary Materials for "*Wolbachia* and host intrinsic reproductive barriers contribute additively to post-mating isolation in spider mites"

### Box S1. Detailed information relating to the populations of *T. urticae* used in this study.

All information concerning the three *T. urticae* populations used in the experiments is provided in Table I. At the time of the experiments, all populations were fully infected with *Wolbachia* [1, 2], and were controlled for the absence of other known bacterial reproductive manipulators (*Cardinium*, *Rickettsia*, *Spiroplasma* and *Arsenophonus*) as described in [3]. Moreover, DNA extractions from pools of 100 females for each population, and subsequent PCR amplification and sequencing of a fragment of the nuclear ribosomal DNA *ITS2* (internal transcribed spacer 2) region, mitochondrial DNA Cytochrome Oxidase subunit I (*COI*), and *Wolbachia wsp* gene were performed using the same procedure and primers as in [3]. Finally, we performed a multilocus sequence typing for *Wolbachia* (MLST; [4]) using primers and protocols described in [2].

We found that the two red populations (Ri1 and Ri2) are strictly identical based on *ITS2* and *COI* sequences, and carry the same *Wolbachia* strain based on the *wsp* gene and MLST profile. However, they differ from the green population (Gi) by 1 SNP in the *ITS2* sequence and 21 SNPs in the *COI* sequence (genetic distance of 0.056 with Kimura 2-parameter; rate of variation gamma distribution; shape parameter = 1). The *Wolbachia* strain infecting the red (Ri1 and Ri2) and the green (Gi) populations also differ based on their MLST profile, with 1 SNP difference in fbpA and coxA sequences (Table I).

**Table I. Spider mite populations and *Wolbachia* infection.**

| Population | | Ri1 | Ri2 | Gi |
|---|---|---|---|---|
| Host information | **Original name** | AMP | LOU | TOM |
| | **Form** | red | red | green |
| | **Collection date** | 18/11/2013 | 03/10/2013 | --/05/2010 |
| | **Host plant** | *Datura stramonium* | *Solanum melongena* | *Solanum lycopersicum* |
| | **Location** | Aldeia da Mata Pequena | Lourinhã | Carregado |
| | **Coordinates** | 38.534363, -9.191163 | 39.248145, -9.276321 | 39.078962, -8.993656 |
| | **Initial number** | 65♀ | 300♀ | 300♀ |
| | ***ITS2*[1]** | GU565314 | GU565314 | AM408031 |
| | ***COI*[1]** | MF428440 | MF428440 | HM486513 |
| *Wolbachia* information | **Isolate (id)[2]** | Turt_B_wUrtAmp (1858) | Turt_B_wUrtAmp (1858) | Turt_B_wUrtTom (1857) |
| | **Strain[2]** | 491 | 491 | 280 |
| | ***gatB* allele[2]** | 9 | 9 | 9 |
| | ***coxA* allele[2]** | 38 | 38 | 164 |
| | ***hcpA* allele[2]** | 143 | 143 | 143 |
| | ***ftsZ* allele[2]** | 23 | 23 | 23 |
| | ***fbpA* allele[2]** | 444 | 444 | 4 |
| | ***wsp*[1]** | GU014541 | GU014541 | GU014541 |
| | **CI level** | 57% | 30% | 0% |
| **References** | | [2, 3] | [2, 3] | [1, 5]; this study |

[1] GenBank accession number matching with 100% coverage and identity at the nucleotide level.
[2] Allele number in the PubMLST *Wolbachia* MLST database, available at http://www.pubmlst.org/wolbachia/

**Box S2. Detailed description of the minor differences in the experimental procedure used to perform the crosses of category 5 in Experiment 1.**

Because simultaneously performing all possible crosses between 3 different populations, infected or not by *Wolbachia*, represents a very important work load for a single experimenter (*i.e.* a total of 36 types of crosses were performed), inter-population crosses using infected females and uninfected males (crosses of category 5; *cf*. Table 1) were performed *ca*. 23 months after the others.

These crosses were initially excluded because they do not inform on *Wolbachia*-induced CI (wCI), and because host incompatibility (HI) was already assessed by performing the crosses of category 3 (*i.e.* inter-population crosses using uninfected females and uninfected males; *cf.* Table 1). Nevertheless, we subsequently decided to perform them, to confirm that *Wolbachia* infection in females does not affect the pattern of HI (*cf.* Table 1).

The general procedure used to perform these crosses was the same as for the other crosses (*cf.* Methods) with some minor differences. First, to reduce workload, the number of mated females used to create age cohorts was 2*100, instead of 3*100 (*cf.* Methods); and males were directly obtained from female age cohorts (*i.e.* no male age cohorts were prepared). Second, due to an overlap between different experiments performed simultaneously in our laboratory, the growth chamber used for the crosses of categories 1-4 was not available, and we had to use a different growth chamber, with no humidity control. As humidity affects development time in spider mites (Suzuki et al. 2012), female age cohorts for these crosses were created 11 days prior to the onset of the experiment (instead of 10 days; *cf.* Methods). Moreover, because humidity is also a main factor affecting egg hatchability in this species [6], it may explain the inflated embryonic mortality rates observed for this category of crosses (*cf.* Figure S1), and overall does not allow for quantitative comparison between the results obtained in these crosses and the previous ones.

**Table S1. Overproduction of males, female embryonic mortality, juvenile mortality, and resulting hybrid production in all intra- and inter-population crosses using *Wolbachia*-infected and uninfected mites.** Mean (± s.e.) relative proportion of F1 female embryonic mortality ($FM_{corr}$), F1 male overproduction ($MD_{corr}$), F1 juvenile mortality ($JM_{corr}$) and production of F1 adult females (FP) in the brood for each type of cross, as well as the total number of replicates excluding females that laid no eggs (N). The $FM_{corr}$ and $JM_{corr}$ indexes remove the basal (embryonic and juvenile, respectively) mortality estimated in control crosses. The $MD_{corr}$ index computes excess production of F1 males relative to the control crosses. Identical or absent superscripts indicate nonsignificant differences between crosses at the 5% level. Values revealing incompatibilities are highlighted by shaded background (green: MD-type; red: FM-type; orange: both types).

| Category | Cross (♀ x ♂) | $MD_{corr}$ (%)[1] | $FM_{corr}$ (%)[2] | $JM_{corr}$ (%)[3] | FP (%)[1] | N[4] |
|---|---|---|---|---|---|---|
| *1* | Ru1 x Ru1 | 14.47 ± 3.86$^{a}$ | 4.01 ± 1.53$^{a}$ | 4.40 ± 1.26 | 50.83 ± 3.41$^{a}$ | 48 |
| | Ru2 x Ru2 | 16.66 ± 5.08$^{a}$ | 5.77 ± 2.17$^{a}$ | 5.65 ± 2.51 | 44.75 ± 3.66$^{a}$ | 47 |
| | Gu x Gu | 18.56 ± 4.61$^{a}$ | 0.19 ± 0.19$^{a}$ | 4.83 ± 1.21 | 46.15 ± 3.56$^{a}$ | 49 |
| | Ri1 x Ri1 | 6.97 ± 2.95$^{a}$ | 2.59 ± 1.03$^{a}$ | 4.49 ± 1.75 | 57.56 ± 2.68$^{a}$ | 50 |
| | Ri2 x Ri2 | 8.18 ± 3.18$^{a}$ | 4.15 ± 1.29$^{a}$ | 3.97 ± 1.27 | 52.27 ± 2.85$^{a}$ | 43 |
| | Gi x Gi | 17.45 ± 4.38$^{a}$ | 1.97 ± 0.60$^{a}$ | 4.82 ± 1.88 | 46.39 ± 4.01$^{a}$ | 49 |
| *2* | Ru1 x Ri1 | 12.43 ± 3.78$^{a}$ | 42.72 ± 4.07$^{c}$ | 1.68 ± 0.56 | 23.45 ± 3.00$^{b}$ | 49 |
| | Ru2 x Ri2 | 21.52 ± 5.58$^{a}$ | 30.88 ± 4.25$^{c}$ | 1.73 ± 0.80 | 27.01 ± 3.29$^{b}$ | 44 |
| | Gu x Gi | 12.64 ± 4.06$^{a}$ | 1.29 ± 0.51$^{a}$ | 2.87 ± 0.85 | 52.81 ± 3.43$^{a}$ | 47 |
| *3* | Ru1 x Ru2 | 19.97 ± 4.35$^{a}$ | 3.78 ± 1.21$^{a}$ | 5.40 ± 1.40 | 45.91 ± 3.39$^{a}$ | 50 |
| | Ru1 x Gu | 17.30 ± 4.20$^{a}$ | 4.84 ± 2.34$^{a}$ | 1.54 ± 0.95 | 50.96 ± 3.48$^{a}$ | 48 |
| | Ru2 x Ru1 | 9.82 ± 3.66$^{a}$ | 2.98 ± 1.39$^{a}$ | 4.50 ± 1.35 | 50.82 ± 3.16$^{a}$ | 41 |
| | Ru2 x Gu | 13.10 ± 4.50$^{a}$ | 3.16 ± 1.05$^{a}$ | 4.92 ± 2.16 | 51.19 ± 3.25$^{a}$ | 44 |
| | Gu x Ru1 | 53.94 ± 4.02$^{b}$ | 10.51 ± 4.52$^{b}$ | 5.71±1.33 | 12.71 ± 2.14$^{c}$ | 47 |
| | Gu x Ru2 | 54.13 ± 4.84$^{b}$ | 2.90 ± 2.02$^{a}$ | 7.46 ± 2.03 | 12.68 ± 2.26$^{c}$ | 49 |
| *4* | Ru1 x Ri2 | 17.14 ± 4.50$^{a}$ | 33.82 ± 4.48$^{c}$ | 2.84 ± 0.80 | 25.27 ± 3.09$^{b}$ | 46 |
| | Ru1 x Gi | 6.78 ± 2.01$^{a}$ | 2.77 ± 1.01$^{a}$ | 3.71 ± 2.11 | 60.53 ± 2.30$^{a}$ | 49 |
| | Ru2 x Ri1 | 10.31 ± 3.96$^{a}$ | 30.25 ± 4.40$^{c}$ | 3.38 ± 0.87 | 27.01 ± 3.29$^{b}$ | 44 |
| | Ru2 x Gi | 12.49 ± 3.69$^{a}$ | 4.32 ± 1.53$^{a}$ | 2.71 ± 1.13 | 49.42 ± 3.49$^{a}$ | 45 |
| | Gu x Ri1 | 46.36 ± 5.06$^{b}$ | 33.31 ± 7.36$^{c}$ | 2.62 ± 0.71 | 8.93 ± 1.99$^{d}$ | 48 |
| | Gu x Ri2 | 58.13 ± 4.49$^{b}$ | 24.57 ± 5.43$^{c}$ | 5.18 ± 1.46 | 7.46 ± 1.66$^{d}$ | 48 |
| | Ri1 x Ri2 | 16.84 ± 4.50$^{a}$ | 2.73 ± 0.86$^{a}$ | 3.43 ± 1.06 | 50.59 ± 3.56$^{a}$ | 49 |
| | Ri1 x Gi | 5.59 ± 2.47$^{a}$ | 5.08 ± 2.04$^{a}$ | 3.41 ± 0.87 | 57.43 ± 2.61$^{a}$ | 47 |
| | Ri2 x Ri1 | 6.45 ± 2.66$^{a}$ | 3.59 ± 1.33$^{a}$ | 1.34 ± 0.56 | 57.51 ± 2.48$^{a}$ | 48 |
| | Ri2 x Gi | 9.99 ± 3.24$^{a}$ | 9.79 ± 2.81$^{b}$ | 2.29 ± 0.71 | 48.53 ± 2.68$^{a}$ | 49 |
| | Gi x Ri1 | 53.73 ± 5.29$^{b}$ | 30.56 ± 4.02$^{c}$ | 2.87 ± 0.99 | 10.64 ± 1.89$^{d}$ | 49 |
| | Gi x Ri2 | 64.25 ± 4.09$^{b}$ | 22.21 ± 5.86$^{c}$ | 2.22 ± 0.69 | 7.90 ± 1.51$^{d}$ | 49 |
| *5* | Ri1 x Ru1 | 12.40 ± 4.22$^{A}$ | 5.34 ± 1.35$^{A}$ | 2.12 ± 0.69 | 52.25 ± 2.92$^{A}$ | 49 |
| | Ri2 x Ru2 | 12.46 ± 3.86$^{A}$ | 5.42 ± 2.24$^{A}$ | 1.85 ± 0.65 | 47.06 ± 3.82$^{B}$ | 45 |
| | Gi x Gu | 30.30 ± 5.55$^{B}$ | 2.02 ± 0.80$^{A}$ | 2.97 ± 1.13 | 39.46 ± 4.28$^{AB}$ | 50 |
| | Ri1 x Ru2 | 9.42 ± 3.85$^{A}$ | 3.96 ± 1.72$^{A}$ | 3.31 ± 0.78 | 54.26 ± 3.24$^{A}$ | 45 |
| | Ri1 x Gu | 8.95 ± 3.41$^{A}$ | 2.16 ± 1.00$^{A}$ | 1.79 ± 0.42 | 57.08 ± 2.55$^{A}$ | 46 |
| | Ri2 x Ru1 | 5.94 ± 3.04$^{A}$ | 8.91 ± 2.49$^{A}$ | 3.55 ± 1.06 | 52.56 ± 2.49$^{B}$ | 46 |
| | Ri2 x Gu | 17.94 ± 4.03$^{B}$ | 6.28 ± 2.16$^{A}$ | 2.89±0.72 | 45.51 ± 3.15$^{B}$ | 48 |
| | Gi x Ru1 | 57.90 ± 4.25$^{C}$ | 9.23 ± 3.42$^{AB}$ | 2.63 ± 0.76 | 10.21 ± 1.81$^{C}$ | 48 |
| | Gi x Ru2 | 64.49 ± 4.66$^{C}$ | 20.15 ± 4.42$^{B}$ | 1.24 ± 0.42 | 9.17 ± 2.14$^{C}$ | 46 |

[1] Includes all crosses for which at least one offspring reached adulthood.
[2] Includes all crosses that produced more than one female offspring.
[3] Includes all crosses for which at least one egg hatched.
[4] Includes all females that laid at least one egg.

**Table S2. F2 offspring production and unviability.** The production and mortality of F2 offspring stemming from F1 virgin females or from F1 males backcrossed with females from their maternal population are displayed for each type of F0 cross. The left part of table provides the number of F1 virgin females that laid at least one egg (#fertile), and the mean (± s.e.) daily oviposition per female over 4 days (#eggs), proportion of unhatched eggs (*i.e.* embryonic mortality; $mEM_{corr}$) and proportion of dead juveniles (*i.e.* juvenile mortality; $mJM_{corr}$), as well as the total number of F1 females tested (N). The right part of the table provides the number of F1 males that sired at least one daughter (#fertile; as only females are sired by males in haplodiploids), and the mean (± s.e.) proportion of females among adult F2 offspring (sex ratio; SR), proportion of unhatched eggs among F2 females (F2 female embryonic mortality, $fEM_{corr}$) and proportion of dead juveniles among F2 females (F2 female juvenile mortality; $fJM_{corr}$), as well as the total number of F1 males tested (N). The $mEM_{corr}$, $fEM_{corr}$, $mJM_{corr}$ and $fJM_{corr}$ indexes, which are estimates of unviability due to hybrid breakdown, remove the basal embryonic and juvenile mortality estimated in control crosses. na: not applicable. F0 crosses in which incompatibilities were found at the F1 are highlighted by shaded background (green: MD-type; red: FM-type; orange: both types). Identical or absent superscripts indicate nonsignificant differences between crosses at the 5% level.

| Parents of tested F1 (F0♀ x F0♂) | Offspring produced by virgin F1 females | | | | | Offspring produced by backcrossed F1 males | | | | |
|---|---|---|---|---|---|---|---|---|---|---|
| | #fertile[1] | #eggs[2] | $mEM_{corr}$ (%)[2] | $mJM_{corr}$ (%)[3] | N[1] | #fertile[1] | $fEM_{corr}$ (%)[4] | $fJM_{corr}$ (%)[4] | SR (%)[4] | N[1] |
| Ru1 x Ru1 | 91 | 5.77 ± 0.24$^{a}$ | 5.38 ± 1.76$^{a}$ | 4.76 ± 1.30$^{ab}$ | 96 | 41 | 8.33 ± 2.36$^{a}$ | 10.62 ± 3.13$^{b}$ | 66.60 ± 2.85$^{a}$ | 49 |
| Gu x Gu | 97 | 5.63 ± 0.17$^{a}$ | 2.86 ± 0.66$^{a}$ | 3.51 ± 0.58$^{ab}$ | 100 | 58 | 4.95 ± 1.32$^{b}$ | 8.75 ± 2.04$^{b}$ | 63.53 ± 2.35$^{b}$ | 85 |
| Ru1 x Ri1 | 81 | 6.95 ± 0.25$^{b}$ | 11.00 ± 2.94$^{a}$ | 2.09 ± 0.90$^{ab}$ | 85 | 68 | 4.23 ± 1.54$^{a}$ | 8.05 ± 1.93$^{b}$ | 67.89 ± 1.36$^{a}$ | 91 |
| Gu x Gi | 96 | 6.31 ± 0.21$^{b}$ | 2.58 ± 0.89$^{a}$ | 6.04 ± 1.57$^{ab}$ | 100 | 58 | 8.18 ± 1.82$^{b}$ | 5.08 ± 1.66$^{a}$ | 64.57 ± 2.46$^{b}$ | 84 |
| Ru1 x Gu | 2 | 0.63 ± 0.13 | 100.00 ± 0.00 | - | 100 | 72 | 3.56 ± 1.39$^{a}$ | 7.12 ± 1.93$^{a}$ | 68.53 ± 1.46$^{a}$ | 84 |
| Gu x Ru1 | 2 | 0.25 ± 0.00 | 100.00 ± 0.00 | - | 96 | 57 | 5.85 ± 1.36$^{b}$ | 5.79 ± 1.63$^{b}$ | 66.24 ± 2.11$^{b}$ | 82 |
| Ru1 x Gi | 0 | - | - | - | 100 | 56 | 2.09 ± 0.65$^{a}$ | 3.00 ± 1.03$^{a}$ | 70.99 ± 1.41$^{a}$ | 69 |
| Gu x Ri1 | 0 | - | - | - | 71 | 45 | 7.87 ± 2.38$^{b}$ | 12.07 ± 2.66$^{b}$ | 54.81 ± 3.24$^{c}$ | 82 |
| Ri1 x Ru1 | 98 | 7.45 ± 0.22$^{c}$ | 5.86 ± 1.33$^{ab}$ | 2.66 ± 0.66$^{a}$ | 100 | 35 | 7.27 ± 2.58 | 7.14 ± 2.55 | 69.34 ± 1.06$^{A}$ | 42 |
| Gi x Gu | 95 | 6.61 ± 0.24$^{b}$ | 4.86 ± 1.01$^{b}$ | 5.30 ± 1.13$^{b}$ | 100 | 26 | 2.36 ± 0.86 | 0.95 ± 0.52 | 69.53 ± 2.41$^{A}$ | 46 |
| Ri1 x Ri1 | 54 | 5.64 ± 0.31$^{a}$ | 2.40 ± 1.46$^{a}$ | 1.83 ± 0.63$^{a}$ | 57 | 10 | 6.49 ± 5.28 | 4.67 ± 3.05 | 69.96 ± 2.90$^{A}$ | 14 |
| Gi x Gi | 75 | 6.34 ± 0.30$^{b}$ | 5.10 ± 1.53$^{b}$ | 5.69 ± 1.22$^{b}$ | 80 | 20 | 3.00 ± 1.12 | 5.54 ± 2.86 | 61.37 ± 4.54$^{B}$ | 37 |
| Ri1 x Gu | 0 | - | - | - | 100 | 25 | 7.67 ± 3.00 | 1.10 ± 0.52 | 65.77 ± 3.54$^{A}$ | 31 |
| Gi x Ru1 | 0 | - | - | - | 100 | 33 | 9.64 ± 2.94 | 6.09 ± 2.77 | 66.53 ± 2.58$^{A}$ | 46 |
| Ri1 x Gi | 1 | 1.25 ± na | 100.00 ± na | - | 100 | 31 | 2.64 ± 0.85 | 4.76 ± 2.18 | 71.64 ± 1.26$^{A}$ | 33 |
| Gi x Ri1 | 1 | 0.75 ± na | 100.00 ± na | - | 93 | 31 | 4.15 ± 1.18 | 4.41 ± 1.39 | 62.50 ± 2.53$^{A}$ | 47 |

[1] All tested individuals

[2] Includes all F1 females that laid at least one egg. Note, given that only a few F1 females resulting from inter-population crosses laid eggs, they were excluded from analyses.

[3] Includes all F1 females that laid at least one egg that hatched

[4] Includes all F1 males that sired at least one daughter

**Table S3. Description of all statistical models used in the experiments.** All response variables were analysed using the glmmTMB procedure, with the type of cross fit as a fixed explanatory variable and the experimental block as random explanatory variable, *i.e.* y ~ cross+(1|block). For the analyses of F1 female/male fertility, the number of days each female/male was alive over the 4-day oviposition period was added to the minimal models as it significantly improved their fit. The "sample size" column gives the number of individual crosses included in each analysis, and the "Family" column indicates the error structure used in each model (bb: betabinomial, zibb: zero-inflated betabinomial, b: binomial, n: log-linked gaussian). Models with (beta)binomial error structure require either a binary response variable (fertile or sterile), a concatenated response variable binding together the number of successes and failures for a given outcome (*e.g.* being a female or not, for female proportion and sex-ratio), or a proportion (bounded between 0 and 1, for all corrected variables). In the latter case, a "weights" argument was added in the model to account for the number of observations per replicate (*i.e.* the denominator). F1 female/male fertility: whether each cross produced at least 1 egg/daughter (1) or none (0); $MD_{obs}$: percentage of adult F1 males relative to total eggs, CCMD: mean percentage of adult F1 males relative to total eggs in control crosses, $FM_{obs}$: percentage of unhatched eggs relative to adult females, CCFM: mean percentage of unhatched eggs relative to females in control crosses, $(m/f)JM_{obs}$: percentage of dead juveniles relative to the total number of eggs (for $JM_{obs}$ and $mJM_{obs}$) or to the total number of females (for $mJM_{obs}$), CC(m/f)JM: mean percentage of dead juveniles relative to the total number of eggs (for $mJM_{obs}$) or to the total number of females (for $fJM_{obs}$) in control crosses, $(m/f)EM_{obs}$: percentage of unhatched eggs relative to the total number of eggs (for $mEM_{obs}$) or to the total number of females (for $fEM_{obs}$), CC(m/f)EM: mean percentage of unhatched eggs relative to the total number of eggs (for CCmEM) or to the total number of females (for CCfEM) in control crosses, daughters/sons: total number of adult daughters/sons produced in each cross, eggs: total number of eggs laid by each female during the oviposition period, day: number of days during which each female used in the experiments was alive.

| | Variable of interest | Response variable | Model No. | Data subset | Sample size | Family | Effect of cross |
|---|---|---|---|---|---|---|---|
| Experiment 1 – F0 crosses | Male overproduction ($MD_{corr}$) | $(MD_{obs}$-CCMD)/(1-CCMD) | 1.1 | Crosses from categories 1-4 | 1263[1] | bb | $\chi^2_{26}$=460.70, $p$<0.0001 |
| | | | 1.2 | Crosses from category 5 | 422[1] | bb | $\chi^2_{8}$=174.26, $p$<0.0001 |
| | Female mortality ($FM_{corr}$) | $(FM_{obs}$-CCFM)/(1-CCFM) | 1.3 | Crosses from categories 1-4 | 918[2] | bb | $\chi^2_{26}$=506.20, $p$<0.0001 |
| | | | 1.4 | Crosses from category 5 | 325[2] | bb | $\chi^2_{8}$=35.85, $p$<0.0001 |
| | Juvenile mortality ($JM_{corr}$) | $(JM_{obs}$-CCJM)/(1-CCJM) | 1.5 | Crosses from categories 1-4 | 1266[3] | bb | $\chi^2_{26}$=34.49, $p$=0.12 |
| | | | 1.6 | Crosses from category 5 | 422[3] | bb | $\chi^2_{8}$=9.13, $p$=0.33 |
| | Female proportion (FP) | cbind(daughters,eggs-daughters) | 1.7 | Crosses from categories 1-4 | 1263[1] | zibb | $\chi^2_{26}$=966.45, $p$<0.0001 |
| | | | 1.8 | Crosses from category 5 | 422[1] | zibb | $\chi^2_{8}$=278.23, $p$<0.0001 |
| Experiment 2 – F1 ♀ | F1 female fertility | Proportion of fertile F1 females | 2.1 | Complete dataset | 1478[4] | b | $\chi^2_{15}$=214.26, $p$<0.0001 |
| | F1 female daily oviposition | eggs/day | 2.2 | F1 females from intra-population crosses | 687[5] | n | $\chi^2_{7}$=55.65, $p$<0.0001 |
| | F2 male embryonic mortality ($mEM_{corr}$) | $(mEM_{obs}$-CCmEM)/(1-CCmEM) | 2.3 | F1 females from intra-population crosses | 687[5] | bb | $\chi^2_{7}$=23.33, $p$=0.001 |
| | F2 male juvenile mortality ($mJM_{corr}$) | $(mJM_{obs}$-CCmJM)/(1-CCmJM) | 2.4 | F1 females from intra-population crosses | 681[3] | bb | $\chi^2_{7}$=18.57, $p$=0.01 |
| Experiment 2 – F1 ♂ | F1 male fertility | Proportion of fertile F1 males | 2.5.1 | Uninfected F1 males | 588[5] | b | $\chi^2_{7}$=25.58, $p$=0.0006 |
| | | | 2.5.2 | Infected F1 males | 276[5] | b | $\chi^2_{7}$=15.23, $p$=0.03 |
| | F2 female embryonic mortality ($fEM_{corr}$) | $(fEM_{obs}$-CCfEM)/(1-CCfEM) | 2.6.1 | Uninfected F1 males | 455[2] | bb | $\chi^2_{7}$=26.31, $p$=0.0004 |
| | | | 2.6.2 | Infected F1 males | 211[2] | bb | $\chi^2_{7}$=5.58, $p$=0.59 |
| | F2 female juvenile mortality ($fJM_{corr}$) | $(JM_{obs}$-CCfJM)/(1-CCfJM) | 2.7.1 | Uninfected F1 males | 455[2] | bb | $\chi^2_{7}$=22.64, $p$=0.002 |
| | | | 2.7.2 | Infected F1 males | 211[2] | zibb | $\chi^2_{7}$=11.68, $p$=0.11 |
| | F2 offspring sex ratio (SR) | cbind(sons, daughters) | 2.8.1 | Uninfected F1 males | 455[2] | bb | $\chi^2_{7}$=42.10, $p$<0.0001 |
| | | | 2.8.2 | Infected F1 males | 211[2] | bb | $\chi^2_{7}$=15.19, $p$=0.03 |

[1] Includes all crosses that produced at least one adult offspring.
[2] Includes all crosses that produced more than one female offspring.
[3] Includes all crosses that produced at least one hatched egg.
[4] Includes all crosses performed in the experiment.
[5] Includes all crosses that produced at least one egg.

**Figure S1. Summary of the development of *T. urticae* eggs resulting from inter-population crosses between infected females and uninfected males (cross category 5).** Bar plots represent mean ± s.e. relative proportions of unhatched eggs (i.e. embryonic mortality), dead juveniles (i.e. juvenile mortality), adult daughters and sons for each type of cross. Mothers are displayed on the bottom level of the x-axis and fathers on the top level.

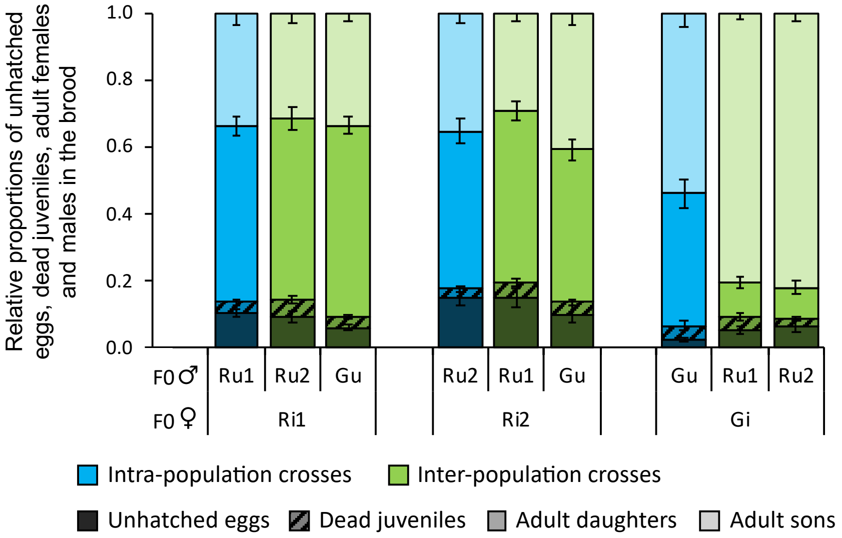

**Figure S2. Overproduction of males, female embryonic mortality, and resulting hybrid production in intra- and inter-population crosses using *Wolbachia*-infected and uninfected mites (cross category 5). (a)** Boxplot of the proportion of males produced in all crosses relative to that in control crosses ($MD_{corr}$). **(b)** Boxplot of the proportion of unhatched eggs relative to females, accounting for the basal level of this proportion observed in control crosses ($FM_{corr}$). **(c)** Proportion of F1 adult females (*i.e.* hybrids) in the brood. Mothers are displayed on the bottom level of the x-axis and fathers on the top level. Identical or absent superscripts indicate nonsignificant differences at the 5% level among crosses.

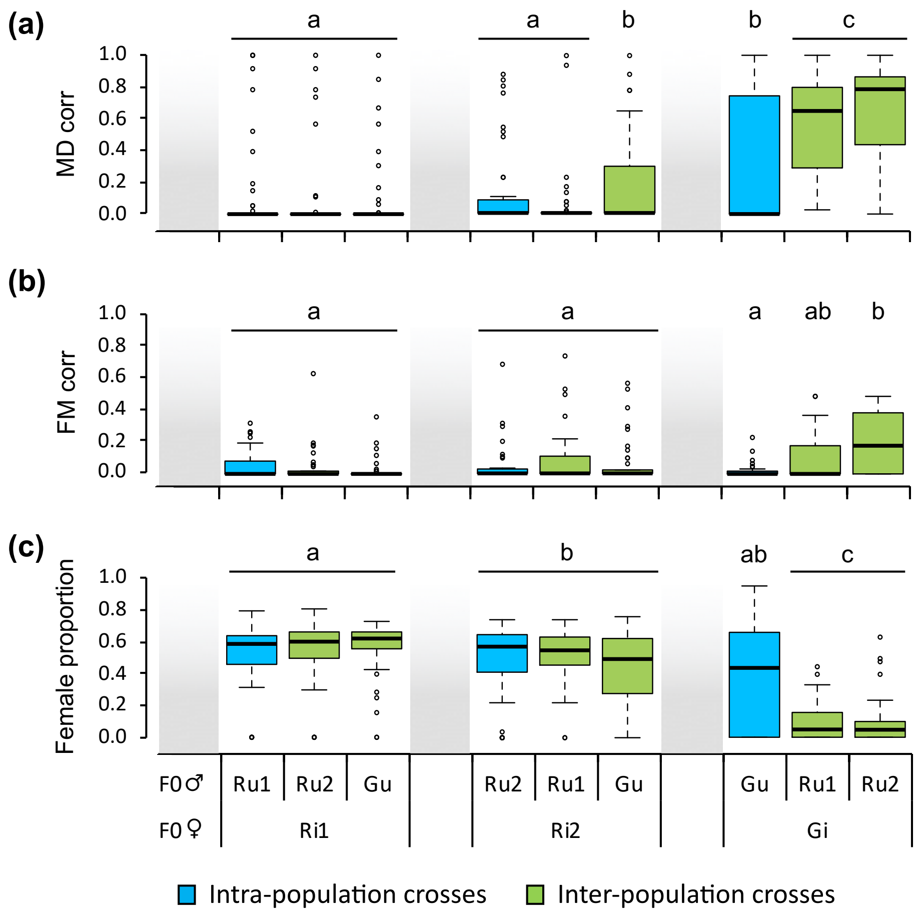

*Note, in* ***(b)*** *some differences in the level of $FM_{corr}$ were found among crosses despite males not carrying Wolbachia (model 1.4, $\chi^2_8$=35.85, p<0.0001). This effect can be attributed to an overestimation of the $FM_{corr}$ parameter when very few daughters were produced due to MD-type incompatibilities, i.e. Gi ♀xRu1 ♂ and Gi ♀xRu2 ♂ crosses. In* ***(a)*** *and* ***(c)****, for an unknown reason, a higher variance was found in the crosses Gi ♀xGu ♂ and Ri2 ♀xGu ♂ than in other crosses not affected by MD-type incompatibilities. Further experiments on mating behaviour are needed to test whether this effect results from a higher proportion of non-mated females in these crosses.*

**Figure S3. Viability of F2 offspring stemming from F1 virgin females.** Boxplots of (a) F2 embryonic mortality estimated using the $mEM_{corr}$ index, and (b) F2 juvenile mortality estimated using the $mJM_{corr}$ index, which accounts for the basal level of mortality (observed in control crosses). The x-axis displays the parents of each tested F1 female. Mothers are displayed on the bottom level of the x-axis and fathers on the top level. Identical or absent superscripts indicate nonsignificant differences at the 5% level among crosses. *Note that no data are displayed for inter-population crosses because all but six F1 females obtained from these crosses were sterile, and none of the few F2 eggs laid by those 6 hybrid females hatched (cf. Table S2).*

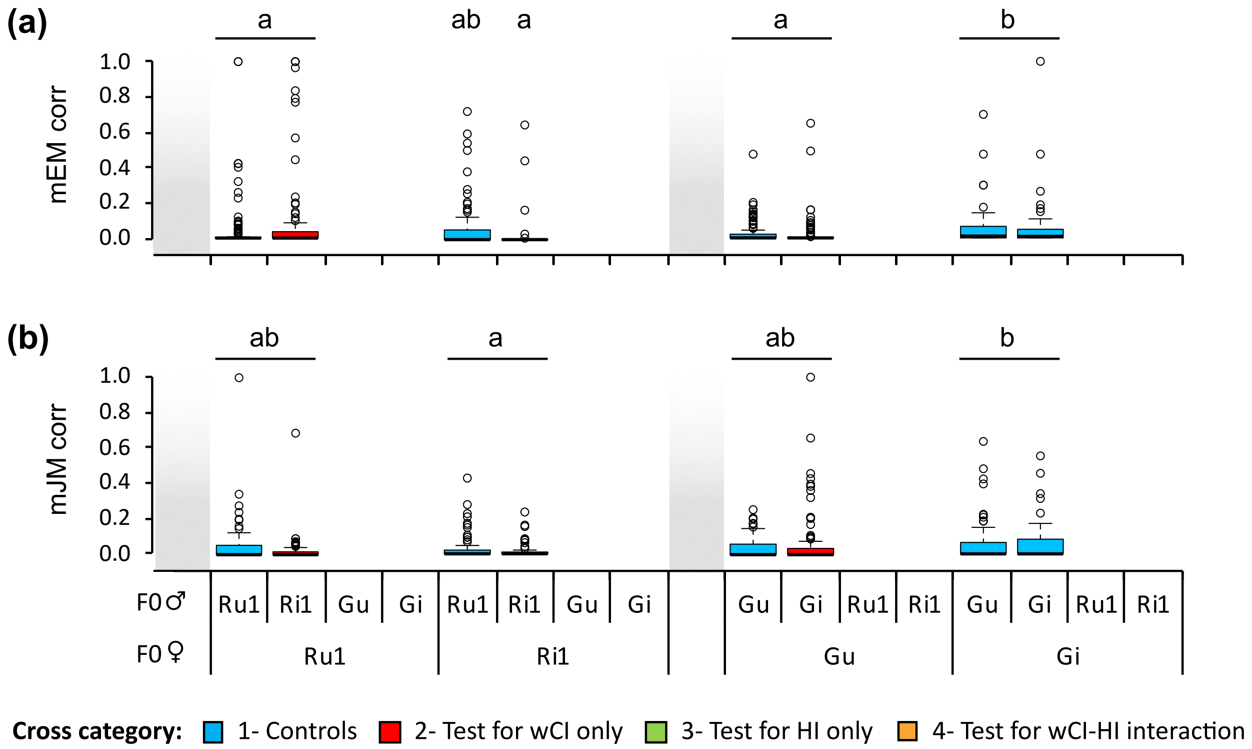

**Figure S4. Viability of F2 offspring stemming from F1 males backcrossed with females from their maternal population.** Boxplots of (a) F2 embryonic mortality estimated using the $fEM_{corr}$ index, and (b) F2 juvenile mortality estimated using the $fJM_{corr}$ index, which accounts for the basal level of mortality (observed in control crosses). The x-axis displays the cross that produced each tested F1 male. F1 males were mated with females from the same population as their mother. Mothers of F1 males are displayed on the bottom level of the x-axis and fathers of F1 males on the top level. Identical or absent superscripts indicate nonsignificant differences at the 5% level among crosses. *Note that crosses using F1 males stemming from uninfected mothers were analysed separately from those using F1 males stemming from infected mothers, as they were performed at different times than all other crosses.*

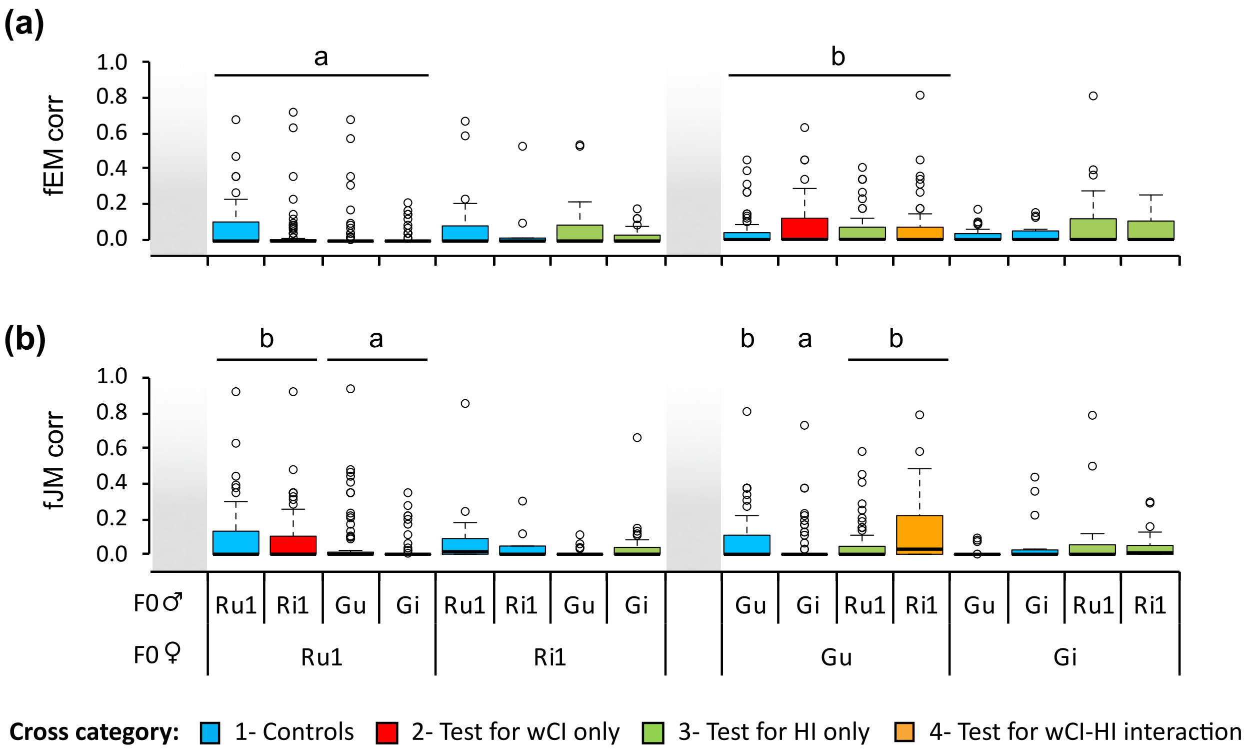
