## Supplementary material for "*Wolbachia* and host intrinsic reproductive barriers contribute additively to post-mating isolation in spider mites": Description of the datasets

Three different datasets are provided (all as Tab delimited Text files):


1. Complete dataset Experiment 1.txt

Dataset used to determine the effects of Wolbachia-induced and host-associated incompatibilities on F1 offspring development. The different cross categories (1 to 5) and cross types (female x male) are given in the columns “category” and “cross”, respectively. The column “block” indicates the temporal block. The column “female_survival” indicates the day at which each female was found dead during the 3-day oviposition period. The column “eggs” indicates the total number of eggs laid. The columns “unhatched” and “hatched” indicate the total number of unhatched and hatched eggs, respectively. The column “dead_juveniles” indicates the total number of offspring which died before reaching adulthood. The columns “daughters” and “sons” indicate the total number of adult female and male offspring, respectively.


2. Complete dataset Experiment 2_Test 1.txt

Dataset used to assess virgin F1 female fertility and viability of their offspring. The column “block” indicates the temporal block. The cross type (female x male) from which the tested females were obtained is provided in the column “cross”. The column “female_survival” indicates the day at which each female was found dead during the 4-day oviposition period. The column “eggs” indicates the total number of eggs laid. The columns “unhatched” and "hatched" indicate the total number of unhatched and hatched eggs, respectively. The column “dead_juveniles” indicates the total number of offspring which died before reaching adulthood, and the column “sons” indicates the total number of adult males produced.


3. Complete dataset Experiment 2_Test 2.txt

Dataset used to assess F1 male fertility when backcrossed to their maternal population, and viability of their offspring. The column “block” indicates the temporal block (U1 to U3 for uninfected males, and I1 and I2 for infected males). The cross type (female x male) from which the tested males were obtained is provided in the column “cross”. The columns “female_survival” and “male_survival” indicate the day at which each female and male, respectively, were found dead during the 4-day oviposition period. The column “eggs” indicates the total number of eggs laid. The columns “unhatched” and “hatched” indicate the total number of unhatched and hatched eggs, respectively. The column “dead_juveniles” indicates the total number of offspring which died before reaching adulthood. The columns “daughters” and “sons” indicate the total number of adult female and male offspring produced, respectively.
